# The role of MYB46 in modulating polysaccharide acetylation by mediating the transcriptional regulation of cell wall-related genes in Arabidopsis

**DOI:** 10.1101/2025.01.01.630977

**Authors:** Lavi Rastogi, Sanjay Deshpande, Prashant Anupama-Mohan Pawar

**Affiliations:** Regional Centre for Biotechnology, NCR, Biotech Science Cluster, Faridabad 121001, India

## Abstract

Understanding the mechanism behind the transcriptional regulation of polysaccharide *O*-acetylation remains a key challenge that might be regulated through transcription factors. Our earlier work revealed the upregulation of *At*GELP7 in MYB46 overexpression lines, prompting us to investigate how MYB46 transcriptionally controls *At*GELP7 including other cell wall acetylation pathway genes in Arabidopsis. In MYB46 overexpression lines, we observed alteration in acetylation levels on xylan, xyloglucan and pectin in different tissue types, which suggests complex and tight regulation of acetylation homeostasis in the cell wall. Further, our transcriptomic data revealed the simultaneous upregulation of both *At*GELP7 (acetyl xylan esterase) and xylan-specific TBLs (xylan *O*-acetyltransferases) indicating sophisticated regulation of cell wall acetylation homeostasis. Using transactivation studies in Nicotiana and *pAtGELP7::GUS* stable lines, we found that MYB46 enhances *At*GELP7 expression probably through downstream regulators which could be either MYB103 or other MYBs. In addition, extensive cell wall analysis of MYB46 overexpression lines showed differential sugar distribution, preferably to major cell wall components such as cellulose, xyloglucan, and xylan across different tissue types of different developmental stages. Moreover, the integration of RNA-sequencing and ChIP-sequencing data uncovered previously unknown, novel probable direct gene targets of MYB46 that may be involved in polysaccharide acetylation and cell wall remodeling.

## Introduction

Plant cell wall biosynthesis and development is a multifactorial process that is well-regulated via multiple transcriptional cues including transcription factors (TFs). Numerous TFs have been identified in the plant kingdom which regulate the expression of genes and thereby control many critical cellular or biological processes. In general, TFs are classified based on the structure of their DNA-binding domain such as basic helix-loop-helix (bHLH), basic region-leucine zipper (bZIP), No apical meristem/*Arabidopsis* transcription activation factor/Cup-shaped cotyledon (NAC), v-myb avian myeloblastosis viral oncogene homolog (MYB), conservative sequence WRKYGOK (WRKY), APETALA2/ethylene-responsive factor (AP2/ERF), etc. Some of them specifically play a significant role in the regulation of cell wall biosynthesis and modifications. Transcriptional regulation of cell wall biosynthesis has been studied through TFs such as NAC and MYBs, which regulate secondary cell wall biosynthesis, whereas AP2/ERFs regulate primary cell wall biosynthesis (Nakano et al., 2015, Sakamoto et al., 2018). The regulation of entire secondary cell wall biosynthesis network is thoroughly investigated and studied through specific master transcriptional regulators such as Secondary wall-associated NAC Domain 1 (SND1) and MYB46 as they act as first-level and second-level main switches of secondary cell wall biosynthesis in Arabidopsis, respectively(Zhong et al., 2006, Zhong et al., 2007). MYB46, being a downstream key regulator, is more likely to directly regulate genes involved in the biosynthesis of major cell wall components, i.e., cellulose, lignin, and hemicellulose, reported previously in the literature (Kim et al., 2013, Kim et al., 2014). Overexpression of these master transcriptional regulators alters the cell wall composition mainly by altering lignin, cellulose, and xylan content. These changes are visualized through lignin-specific phloroglucinol-HCl staining and immunolocalization studies using cellulose and xylan-specific antibodies (Zhong et al., 2006, Zhong et al., 2007). In some reports, crystalline cellulose, lignin, and hemicellulosic content are also quantified in MYB46 overexpressor lines (Kim et al., 2013, Kim et al., 2014). Since the focus of these studies was to understand secondary cell wall biosynthesis, therefore there lies some research gaps like how this increase in cellulose, lignin, and xylan content affects the composition of other cell wall monosugars; Is there any metabolic shifting in such cases to maintain the overall sugar homeostasis? How the cell wall composition differs in different plant tissues and also at different developmental stages. However, the detailed characterization of cell wall components is what is lacking. While researchers have extensively studied how overall secondary cell wall biosynthesis is regulated however, how MYB46 regulates side groups that present on polysaccharide backbone, such as acetyl substitutions, is relatively less explored. Moreover, it is unknown whether MYB46 regulates genes involved in cell wall remodeling and integrity. Additionally, the direct targets involved in these processes remain largely unidentified. Plant cell wall polysaccharides like hemicelluloses (xylan and xyloglucan) and pectin have modifications of *O*-acetyl groups in their backbone and side chain sugar moieties. The major genes/ or proteins that are involved in the polysaccharide acetylation pathway in plants belong to four families i.e., REDUCED WALL *O*-ACETYLATION (RWA), ALTERED XYLOGLUCAN 9 (AXY9), TRICHOME-BIREFRINGENCE-LIKE (TBL), and GDSL ESTERASE/LIPASE PROTEIN (GELP) which are acetyl transporters, acetyl transferases and acetyl esterases, respectively (Pauly and Ramirez, 2018, Rastogi et al., 2022, Rastogi et al., 2024). The transcriptional regulation of these genes involved in cell wall polysaccharide acetylation pathway is scarcely understood. Some previous reports demonstrated the regulation of some of these genes by master transcriptional regulators such as SND1, Vascular-related NAC Domain 7 (VND7) and MYB46. The reports by (Lee et al., 2011, Yuan et al., 2013) showed the upregulation of *RWA1*, *RWA3*, *RWA4*, and *ESK1*/*TBL29* genes in SND1 overexpression lines in Arabidopsis. Cytosolic acetyl-CoA pool, which serves as a probable acetyl donor for the polysaccharide acetylation pathway, is known to be generated by ATP-CITRATE LYASE (ACL) encoded by three *ACLA* (*ACLA-1*, *ACLA-2*, and *ACLA-3*) and two *ACLB* (*ACLB-1* and *ACLB-2*) genes. These genes were also found to be highly induced upon overexpression of SND1, VND7 and MYB46 (Zhong et al., 2020). This shows the positive regulation of polysaccharide acetylation pathway by these transcriptional regulators. A recent study showed the role of a bHLH transcription factor i.e., MYC2 in cell wall acetylation regulation. As a central regulator of jasmonate signalling, MYC2 activates multiple TBLs, particularly *TBL37* and *ESK1*/*TBL29* (Sun et al., 2020). This activation leads to enhanced cell wall acetylation, ultimately resulting in enhanced resistance against herbivores. Also, a recent findings have demonstrated that MYB103 positively regulates TBL27 that controls xyloglucan acetylation and plays a crucial role in plant’s response in aluminium (Al) stress (Wu et al., 2022). While previous studies have focused on genes regulating polysaccharide acetylation in the Golgi, our recent work has identified two novel proteins from Arabidopsis GELP family i.e., *At*GELP7 and *At*GELP53 that localise to the plasma membrane. These proteins functions as acetyl xylan esterase and xyloglucan acetyl esterase, respectively and are involved in modifying polysaccharide acetylation post synthesis (Rastogi et al., 2022, Rastogi et al., 2024). Notably, in plants overexpressing MYB46, the expression of *AtGELP7* gene was higher by 55-fold as compared to wild type. This suggests that MYB46 is involved in regulation of *AtGELP7*. Based on this, the precise mechanism and control of MYB46 in cell wall acetylation, remodeling and integrity needs to be investigated for better understanding of homeostasis mechanism upon disruption of cell wall during growth or defense. In this study, we used MYB46 overexpressing Arabidopsis stable lines for extensive compositional characterization of cell wall components across different tissue types of different developmental stages. In addition, we did cell wall acetylation profiling in all these tissues to examine total cell wall acetyl content and acetylation level on individual polysaccharides. Through transactivation analysis, we specifically found that MYB46 positively regulates *AtGELP7* gene expression but indirectly via other unknown downstream regulator. We further performed RNA sequencing and Chip sequencing analysis of MYB46; combining both these methods, we identified novel targets of MYB46, which are involved in polysaccharide acetylation, remodelling and integrity.

## Results

### *At*MYB46 overexpressing lines show ectopic deposition of cell wall components and localization in the nucleus

To study the transcriptional regulation of cell wall by Arabidopsis MYB46 (*AT5G12870*) transcription factor, we generated transgenic lines overexpressing *AtMYB46* under *35S* constitutive promoter. These plants were represented in our previous study, where we found 55-fold higher expression of *AtGELP7*, an acetyl xylan esterase, upon MYB46 overexpression (Rastogi et al., 2022). In the current study, we further characterized these transgenic lines (*At*MYB46 OE-2 and *At*MYB46 OE-3) for showing elevated expression level of secondary cell wall-related genes and having an ectopic secondary cell wall deposition as reported in (Ko et al., 2009). The transgenic lines overexpressing *35S::AtMYB46* did not show any growth phenotype and morphologically looked similar to wild-type Arabidopsis plants (Rastogi et al., 2022). MYB46 primarily regulates lignin biosynthesis genes along with other cell wall related genes (Kim et al., 2014), therefore, we tested the expression of *PHENYLALANINE AMMONIA LYASE* (*PAL2*), *FERULATE 5-HYDROXYLASE* (*F5H*), and *CINNAMATE 4-HYDROXYLASE* (*C4H*) and their expression was found enhanced in *At*MYB46 OE lines (Figure 1a). ATP-citrate lyase (ACL) downregulation affects cell wall acetylation and the expression of ACLA was upregulated in MYB46 overexpressor lines (Zhong et al., 2020). We also found enhanced expression of multiple *ACLA-1*, *ACLA-2*, and *ACLA-3* in *At*MYB46 OE lines (Figure 1b). Further, we performed histological analysis in main stem transverse sections of *At*MYB46 OE lines to study ectopic deposition of secondary cell wall components. The thickening of cell wall of xylem cells was observed in both *At*MYB46 OE lines as compared to wild-type plants upon toluidine blue O staining (Figure 1c) and phloroglucinol-HCl staining (Figure 1d). Moreover, we also confirmed *At*MYB46 localization in the nucleus by expressing GFP-*At*MYB46 construct in *Nicotiana benthamiana* using transient expression (Figure 1e). We validated these results by also isolating protoplasts from the stable Arabidopsis GFP-*At*MYB46 lines and MYB46 GFP signals colocalized with DAPI stained nucleus (Figure 1f). Collectively, all of these results confirmed our *At*MYB46 overexpressing transgenic lines for further cell wall-related studies.

**Figure 1.**
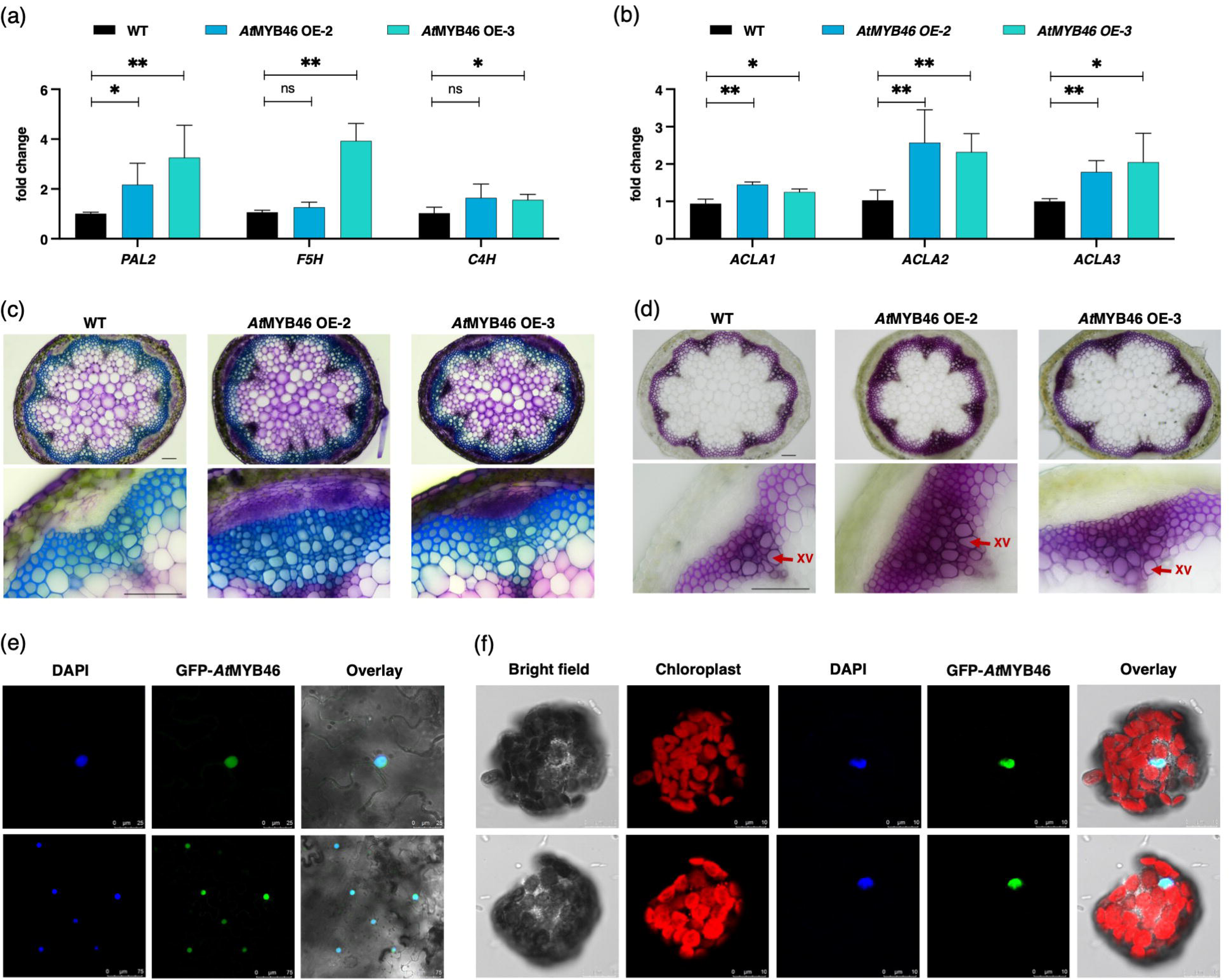
Overexpression of *At*MYB46 shows ectopic cell wall deposition and nuclear localization. The RNA level expression of lignin biosynthetic genes (a), and cell wall acetylation genes (b) were checked in the main stem tissue of *35S::AtMYB46* overexpressing lines by qRT-PCR. Transverse sections of main stems of *35S::AtMYB46* overexpressing lines stained with toluidine blue O (c), and phloroglucinol-HCl (d). Scale bar: 100 µm. (e) Nicotiana leaves were infiltrated with GFP-tagged *At*MYB46 and nuclei stain, i.e., DAPI (4’,6-diamidino-2-phenylindole, dilactate) and visualised under the confocal microscope on 3^rd^ day post infiltration. (f) Visualization of protoplasts isolated from rosette leaves of Arabidopsis stable line of *At*MYB46-GFP under confocal microscope. DAPI was used for nuclei staining. Chloroplast autofluorescence was observed in red channel. Data represents mean ± SE, *n* = 3-4 biological replicates, Student’s t-test at \*\*\*\**p* ≤ 0.001, \*\*\**p* ≤ 0.01, \*\**p* ≤ 0.05, * *p* ≤ 0.1.

### Cell wall sugar distribution in *At*MYB46 overexpression lines shows preferential allocation to major cell wall components

Previous studies by (Kim et al., 2013, Kim et al., 2014) mainly focused on understanding the transcriptional regulation of secondary cell wall biosynthesis; therefore, in such studies, the characterization of cell wall components was limited. In the present study, we investigated alteration in the cell wall composition such as lignin, cellulose, matrix polysaccharides (hemicelluloses and pectin) at different plant developmental stages upon MYB46 overexpression in detail. Therefore, we characterized the cell wall composition in *At*MYB46 OE lines in leaf and stem tissues at different developmental stages. We selected three-week-old rosette leaves, five-week-old cauline leaves, six-seven-week-old dried rosette leaves, five-week-old side stems, main stems from the bottom (10 cm), main stems from the top (10 cm), and six-seven-week-old dried main stems for the analysis. In the leaf tissue, the lignin content was comparable in wild type and *At*MYB46-OE lines whereas the lignin content was increased in all stem tissues of *At*MYB46 OE lines, mainly in the main stem (bottom and top) (Figure 2a, b). Cellulose content was enhanced in six-seven-week-old dried leaf and stem tissues of *At*MYB46 overexpressing lines (Figure 2c, d). Matrix polysaccharides were analyzed by ion chromatography. In leaf tissues, we found a significantly higher content of hemicellulosic and non-crystalline glucose with an increasing percentage in rosette leaf (4%), cauline leaf (10%), and dried rosette leaf (12%) whereas all other monosugars like fucose, rhamnose, arabinose, galactose, mannose, xylose, and galacturonic acid were comparably downregulated in *At*MYB46 OE lines (Figure 2e, f and S1a). In stem tissues, hemicellulosic and non-crystalline glucose was enhanced in side stem, main stem (bottom), and main stem (top), xylose was increased in main stem (bottom) and dried main stem whereas other monosugars showed a similar reducing trend like leaf tissues in transgenic lines (Figure 2g, h and S1b, c). Overall, this data showed that MYB46 overexpression lines exhibited a shift in sugar distribution, with increased allocation towards cellulose, hemicellulosic glucose (xyloglucan), and xylan which is accompanied by reduced levels of other sugars. This probably suggests either change in the expression of genes involved in sugar substitution, such as glycosyl transferase, or altered UDP sugar metabolism (which are possible substrates for polysaccharide elongation and substitution).

**Figure 2.**
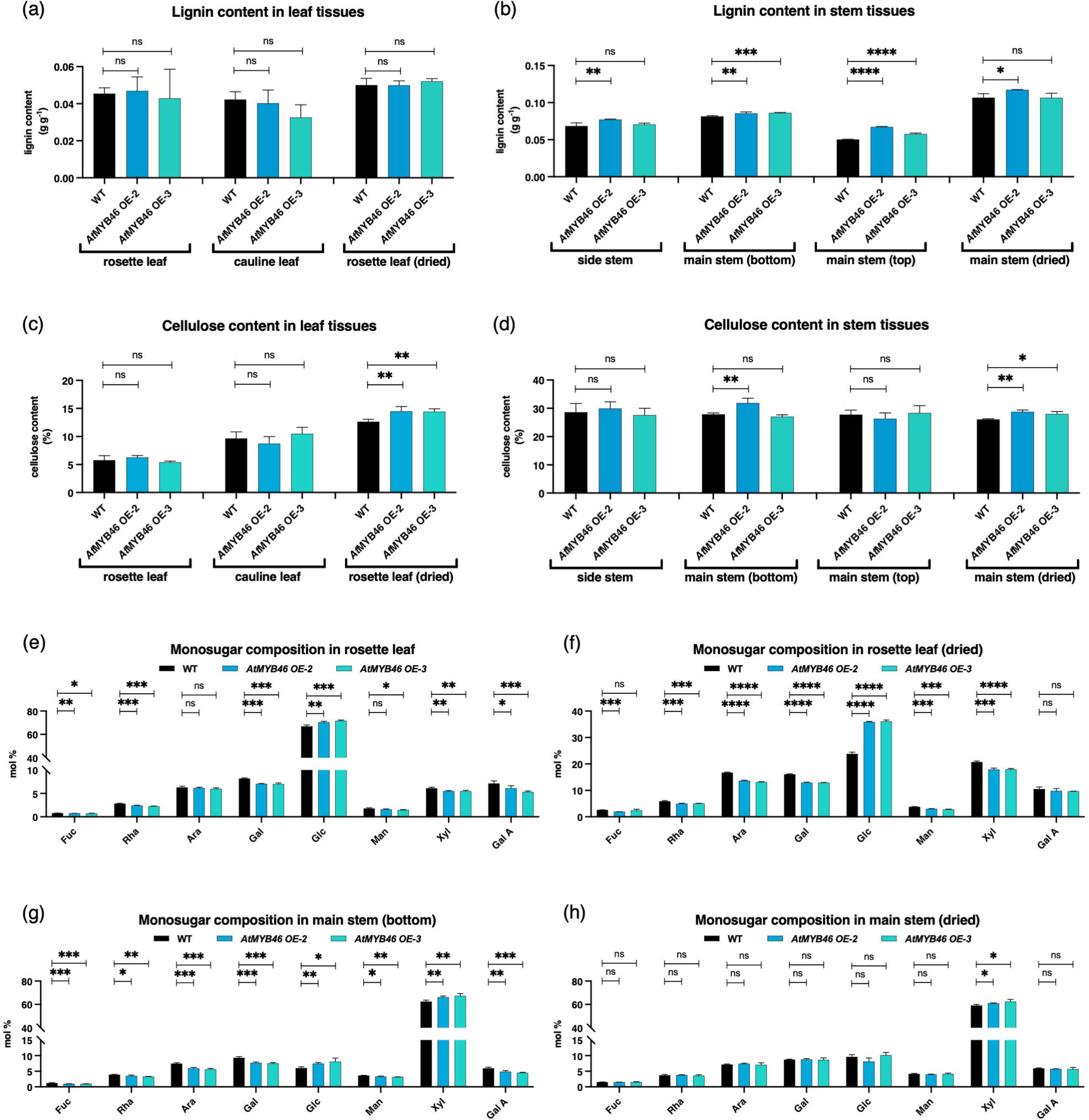
Cell wall compositional characterization of *At*MYB46 overexpression lines reveals differential sugar distribution in different tissue types. Alcohol insoluble residue (AIR) from different plant tissues such as rosette leaf, cauline leaf, rosette leaf (dried), side stem, main stem (bottom), main stem (top), and main stem (dried) of wild-type and *At*MYB46 overexpression (OE) lines were prepared. Lignin content was analyzed in leaf tissues (a), and stem tissues (b) by acetyl bromide soluble lignin (ABSL) method. Cellulose content was detected in leaf tissues (c), and stem tissues (d) by Updegraff method. Monosugar composition analysis (mol%) in leaf tissues (e,f), and stem tissues (g,h) analyzed by ion-chromatography (IC). Fuc-fucose, Rha-rhamnose, Ara-arabinose, Gal-galactose, Glc-glucose, Man-mannose, Xyl-xylose, Gal A-galacturonic acid. Data represents mean ± SE, *n* = 3-4 biological replicates, Student’s t-test at \*\*\*\**p* ≤ 0.001, \*\*\**p* ≤ 0.01, \*\**p* ≤ 0.05, * *p* ≤ 0.1.

### Cell wall polysaccharide *O*-acetylation profiling shows the modulation in individual polysaccharide acetylation levels across different tissues in *At*MYB46 overexpression lines

Transcriptional regulation of gene involved in polysaccharide acetylation is scarcely understood. Some previous reports demonstrated MYB46 and SND1-dependent activation of cell wall acetylation genes such as *RWA1*, *RWA3*, *RWA4*, *ESK1/TBL29*, *AtGELP7*, *ACLA-1*, *ACLA-2*, *ACLA-3*, *ACLB-1* and *ACLB-2* (Lee et al., 2011, Yuan et al., 2013, Zhong et al., 2020, Rastogi et al., 2022). However, whether the disruption of these genes and their effects on the polysaccharide acetyl content level *in planta* was not known. Therefore, we tested total cell wall acetylation levels in different leaf and stem tissues at different developmental stages in wild-type Arabidopsis plants. In the leaf tissues, total cell wall acetyl content level was 5.8 mg/g, 7.5 mg/g, 8 mg/g of AIR in rosette leaf, cauline leaf, dried rosette leaf, respectively whereas in the stem tissues, 27.5 mg/g, 27 mg/g, 24 mg/g, and 25 mg/g of AIR in side stem, bottom main stem, top main stem, and dried main stem, respectively (Figure S2). Collectively, we found that stem tissues have high acetyl content levels in the cell wall, with an average of 26 mg/g of AIR, compared to leaf tissues with an average of 7 mg/g of AIR. Further, we examined the effect of MYB46 overexpression on cell wall acetyl content in different tissue types. A slight reduction was observed in total cell wall acetyl content level in the rosette and cauline leaf tissues in *At*MYB46 OE lines, whereas stem tissues did not show much changes in the acetylation level (Figure 3a, b). To see further changes in acetylation levels in individual polysaccharides upon MYB46 overexpression, we sequentially extracted acetylated polysaccharides i.e., pectin, xyloglucan, and xylan in different fractions from cell wall AIR and acetyl content was analyzed in each fraction. These fractions were also run on ion chromatography to estimate monosaccharide composition in each fraction (Table S1a, b, c). The pectin-rich fraction I extracted in ammonium formate showed slightly increased acetyl content in the rosette leaf, but no significant change was observed in the main stem bottom and dried main stem tissue in *At*MYB46 OE lines (Figure 3c). The pectin-rich fraction II collected after pectate lyase digestion showed an increase in acetyl content in rosette leaf, main stem bottom and dried main stem tissue in *At*MYB46 OE lines compared to wild-type (Figure 3d). The depectinized pellet was digested with xyloglucanase and supernatant containing xyloglucan-rich fraction showed increased acetyl content in rosette leaf but no change was observed in main stem bottom and dried main stem tissue in *At*MYB46 OE lines (Figure 3e). The remaining pellet rich in xylan showed a reduction in acetyl content in rosette leaf and dried main stem tissue from *At*MYB46 OE lines (Figure 3f). This alteration in cell wall acetyl content in pectin, xyloglucan, and xylan represents a tight and complex regulation of cell wall acetylation in different tissue types upon MYB46 overexpression. These changes may be driven by direct or indirect targets of MYB46 which are involved in polysaccharide acetylation including *At*GELP7.

**Figure 3.**
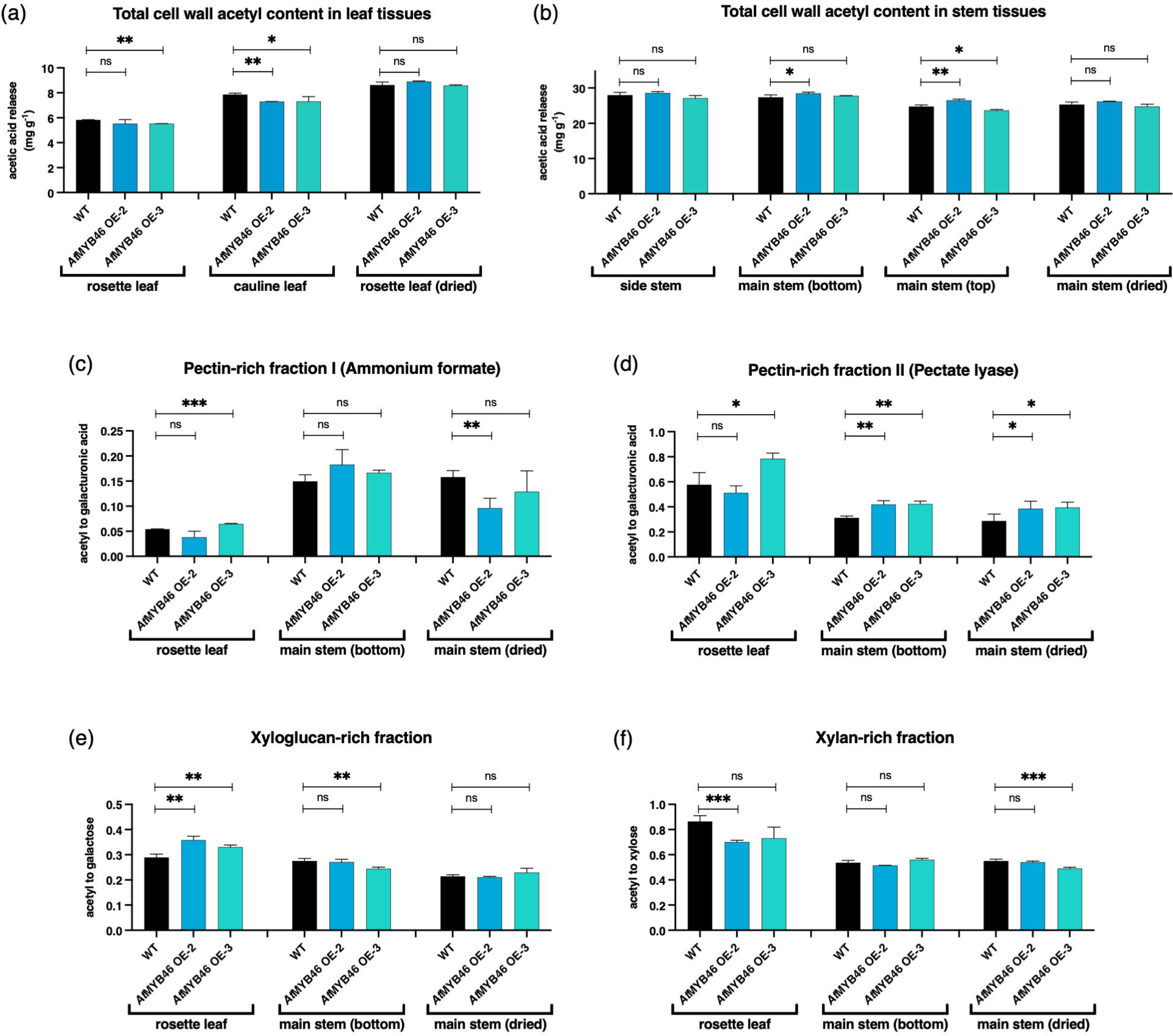
Cell wall polysaccharide *O*-acetylation profiling exhibits tissue-specific modulation in acetylation levels in *At*MYB46 overexpression lines. The total cell wall acetyl content in AIR of different leaf tissues (a) and stem tissues (b) was detected by acetic acid kit after de-esterification. AIR of rosette leaf, main stem (bottom), and main stem (dried) of wild-type and *At*MYB46 OE lines were analyzed for acetyl content in sequentially extracted polysaccharide fractions i.e., pectin-rich fraction I (c), pectin-rich fraction II (d), xyloglucan-rich fraction (e), and xylan-rich fraction (f), and normalized to respective sugar content in each fraction. Data represents mean ± SE, *n* = 3-4 biological replicates, Student’s t-test at \*\*\*\**p* ≤ 0.001, \*\*\**p* ≤ 0.01, \*\**p* ≤ 0.05, * *p* ≤ 0.1.

### Transactivation analysis showed that *At*GELP7 is positively regulated by MYB46 via unknown downstream regulators

We found that a plasma-membrane localized acetyl xylan esterase *At*AXE1/*At*GELP7 was 55-fold upregulated in MYB46 overexpressing lines, suggesting that *At*GELP7 could be the direct or indirect target of MYB46. To investigate this in more detail, we initially measured specific esterase activity using 4-nitrophenyl acetate as substrate in different tissues of MYB46 OE lines and found slightly increased esterase activity in cauline leaf and bottom main stem tissue whereas no change was observed in other tissues (Figure S3a, b). We further checked the expression of *At*GELP7 in different tissues of MYB46 OE lines and found upregulation of *At*GELP7 in all the tissues, as previously observed in (Rastogi et al., 2022) (Figure 4a, b). MYB46 transcription factor recognizes and binds to Secondary cell wall MYB Responsive Elements (SMRE) in their target gene promoter (Zhong and Ye, 2012). Therefore, we looked for SMRE sites in promoter of GELP family gene members belonging to clade Id as shown in (Rastogi et al., 2022). The promoter of *AtGELP7* gene has two SMRE sites i.e., SMRE4 [ACCAACC (-)] and SMRE5 [ACCTAAT (+)] whereas other GELPs from clade Id also have SMRE sites but their expression was not changed upon MYB46 overexpression as observed in (Rastogi et al., 2022) (Table S2). To understand the *AtGELP7* gene regulation by MYB46, we performed transactivation using the effector construct i.e., *35S::AtMYB46* (Figure S4a). The reporter constructs i.e., *pAtGELP7::GUS/LUC*, were made by fusing promoter of *AtGELP7* gene with glucuronidase (GUS) and luciferase (LUC) (Figure S4b). Variable length of promoter was used such as full length *AtGELP7* gene promoter (1.6 kb), promoter region having SMRE5 in forward strand (FS_201 bp), and promoter region having SMRE4 in reverse strand (RS_207bp). The GUS expression was visible, containing only promoter-GUS fusion 1.6Kb and RS_207 promoter (Figure S4c), but GUS expression was not found in FS_201 promoter GUS fusion. Similarly, luciferase was expressed transiently when promoter luciferase was expressed in Nicotiana leaf (Figure S4d). However, GUS and luciferase did not get activated when co-infiltrated with effector construct in Nicotiana leaves (Figure S4c and S4d). These transactivation assays suggested that MYB46 probably is not directly binding to *AtGELP7* gene promoter. However, the expression of *AtGELP7* was upregulated upon MYB46 overexpression, it is possible that there are some downstream regulators through which MYB46 is regulating *AtGELP7*. To understand this further, we generated GUS stable line of *At*GELP7 gene promoter and its expression was detected by GUS staining of different plant parts such as seedlings, leaves, stems, inflorescence, and siliques (Figure 4c). More specifically, *At*GELP7 expression was found higher in stem tissue in comparison to leaf tissue. Therefore, in the leaves of three independent GUS stable lines of *At*GELP7 promoter, we infiltrated the effector construct i.e., *35S::AtMYB46* and further did GUS staining. The GUS expression was enhanced in the leaves infiltrated with MYB46 construct in all the independent lines and this was also quantified (Figure 4d, e). This suggests that MYB46 is regulating *AtGELP7* gene expression, not by binding to its promoter directly but through unknown downstream regulators of Arabidopsis.

**Figure 4.**
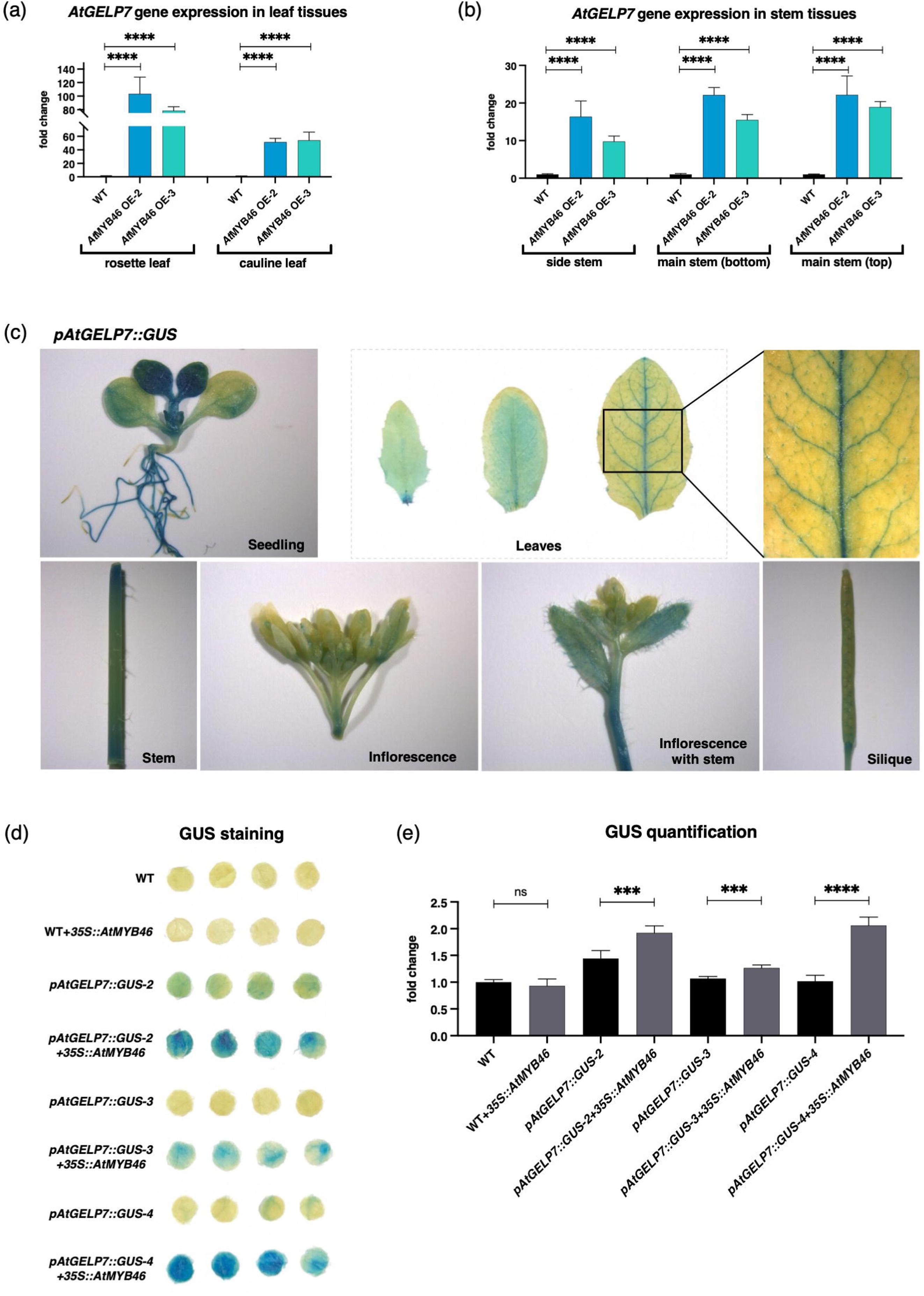
Transactivation study displays regulation of *At*GELP7 by *At*MYB46 transcription factor. The expression of *AtGELP7* gene in different leaf tissues (a), and stem tissues (b) of wild-type and *At*MYB46 OE lines by qRT-PCR. (c) *AtGELP7* gene expression was detected in seedlings, leaves, stems, inflorescence stems, and siliques of *At*GELP7 promoter-GUS stable transgenic line (*pAtGELP7::GUS*) by promoter-GUS activity using X-Gluc as substrate. (d) The rosette leaves of *pAtGELP7::GUS* stable lines were infiltrated with *35S::AtMYB46* Agrobacterium construct and promoter-GUS activity was detected by GUS staining using X-Gluc as substrate. (e) The graph shows GUS quantification represented in fold change. Data represents mean ± SE, *n* = 3-4 biological replicates, Student’s t-test at \*\*\*\**p* ≤ 0.001, \*\*\**p* ≤ 0.01, \*\**p* ≤ 0.05, * *p* ≤ 0.1.

### MYB46 overexpression activates genes belonging to polysaccharide acetylation, biosynthesis and remodeling

To further study the regulation of cell wall acetylation by MYB46, we performed RNA-sequencing analysis on inflorescence stem in wild-type and *At*MYB46 OE-3 line. The principal component analysis (PCA) showed a clear separation of wild-type and *At*MYB46 OE-3 line into two groups with PC1 contributing to 80.3% of variance (Figure 5a). Further analysis revealed 968 differentially expressed genes in MYB46 overexpressing lines with absolute log2 fold change ≥ 1. Of these, 748 genes were significantly upregulated, and 220 genes were significantly downregulated (Figure 5b). Importantly, *AtGELP7* gene expression was to be highest among upregulated genes of *At*MYB46 OE-3 line consistent with our previous findings. The heatmap representation of top 30 differentially expressed genes (DEGs) found to be mainly associated with cell wall biosynthesis and modification in *At*MYB46 OE-3 line (Figure S5). Pathway enrichment analysis of DEGs by Kyoto Encyclopedia of Genes and Genomes (KEGG) revealed over-represented genes involved in various metabolic processes, such as biosynthesis of secondary metabolites, phenylpropanoid biosynthesis, flavonoid biosynthesis, carotenoid biosynthesis, glucosinolate biosynthesis, galactose, starch and sucrose metabolism (Figure 5c).

**Figure 5.**
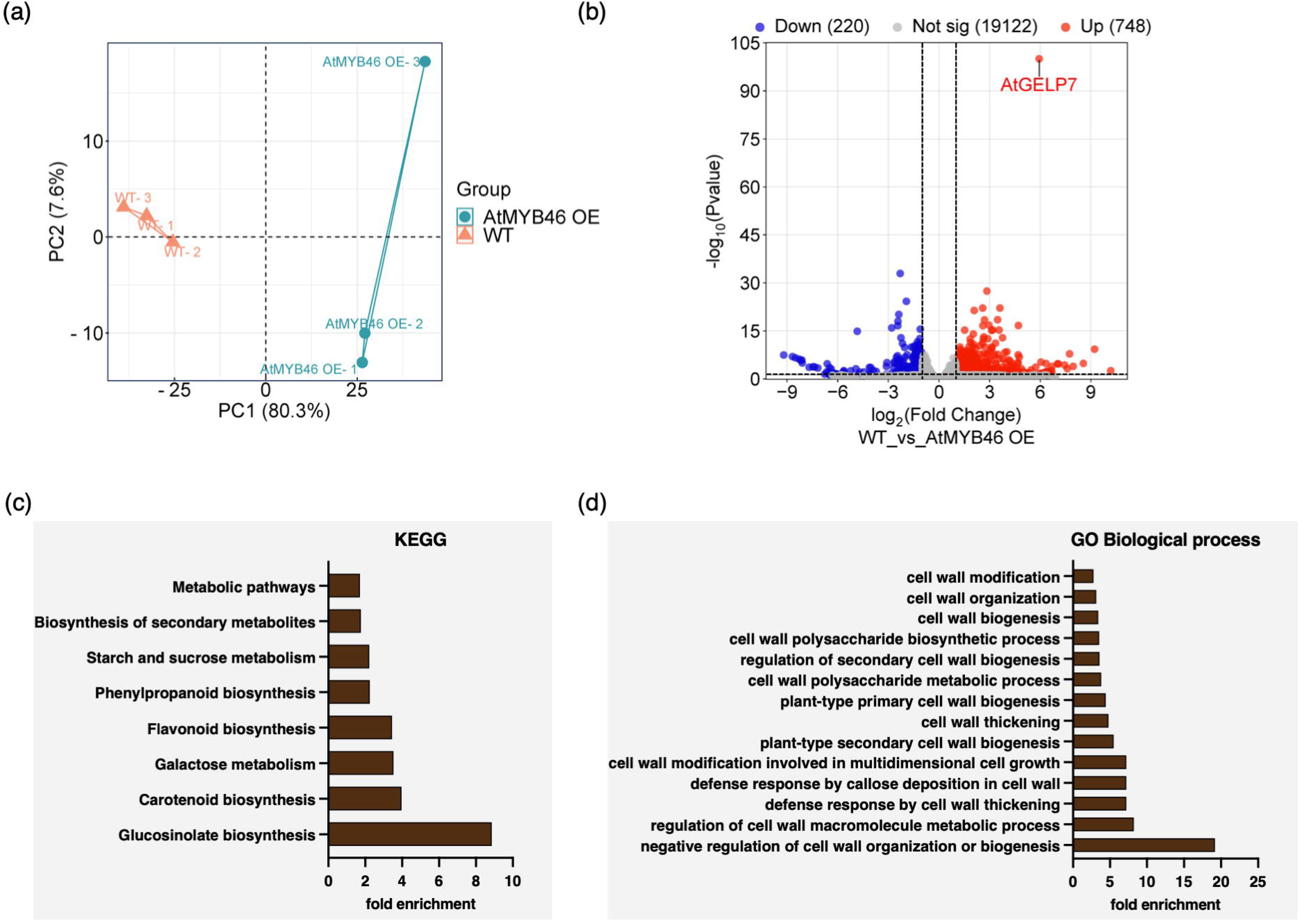
RNA-sequencing of *At*MYB46 overexpression lines shows differential transcriptomic profiles. (a) The principal component analysis (PCA) plot shows a clear distinction between three biological replicates of *At*MYB46 OE main stem samples and the wild-type samples. (b) The volcano plot represents the number of differentially expressed genes (DEGs) i.e., upregulated genes (748) and downregulated genes (220). Pathway analysis of DEGs in *At*MYB46 OE plants by KEGG (c), and Gene Ontology (GO); biological process (d).

The gene ontology (GO) terms represented the involvement of these DEGs in biological processes associated with cell wall thickening, biogenesis, organization, and modification (Figure 5d). The role of MYB46 in secondary cell wall biosynthesis is well understood but poorly in polysaccharide acetylation and cell wall modification. Therefore, we focused on the identification of differentially expressed genes in cell wall acetylation, remodeling and integrity (Table 1). We found 16 differentially expressed cell wall acetylation genes which included *AtGELPs*, *TBLs*, and *ACL*. Most of the *At*GELPs listed in Table 1 are uncharacterized except *At*GELP7, which functions as acetyl xylan esterase (Rastogi et al., 2022). TBLs are polysaccharide acetyltransferases, and the identified TBLs i.e., *ESK1/TBL29*, *TBL33*, and *TBL34* are xylan-specific acetyltransferases whereas *TBL36*, *TBL37*, and *TBL45* are probably involved in esterification of pectin and hemicelluloses (Yuan et al., 2013, Yuan et al., 2016a, Yuan et al., 2016b). ATP-citrate lyase (ACL) genes, i.e., *ACLA-2* and *ACLB-2,* are important for maintaining the cytosolic pool of acetyl CoA, which is a probable acetyl donor for the acetylation pathway (Zhong et al., 2020). Upregulation of these genes suggests the active role of MYB46 in maintaining acetylation homeostasis.

**Table 1.**
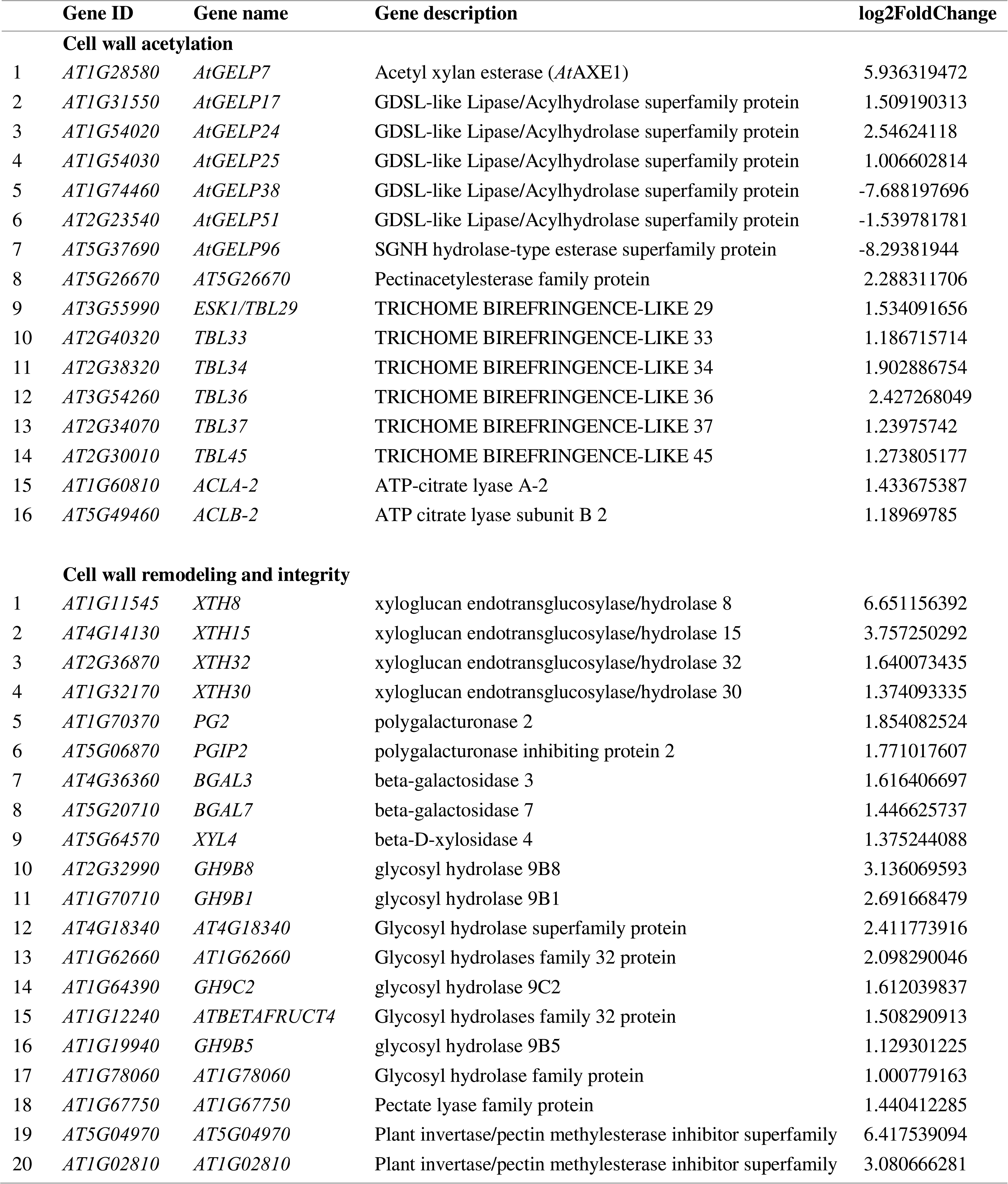
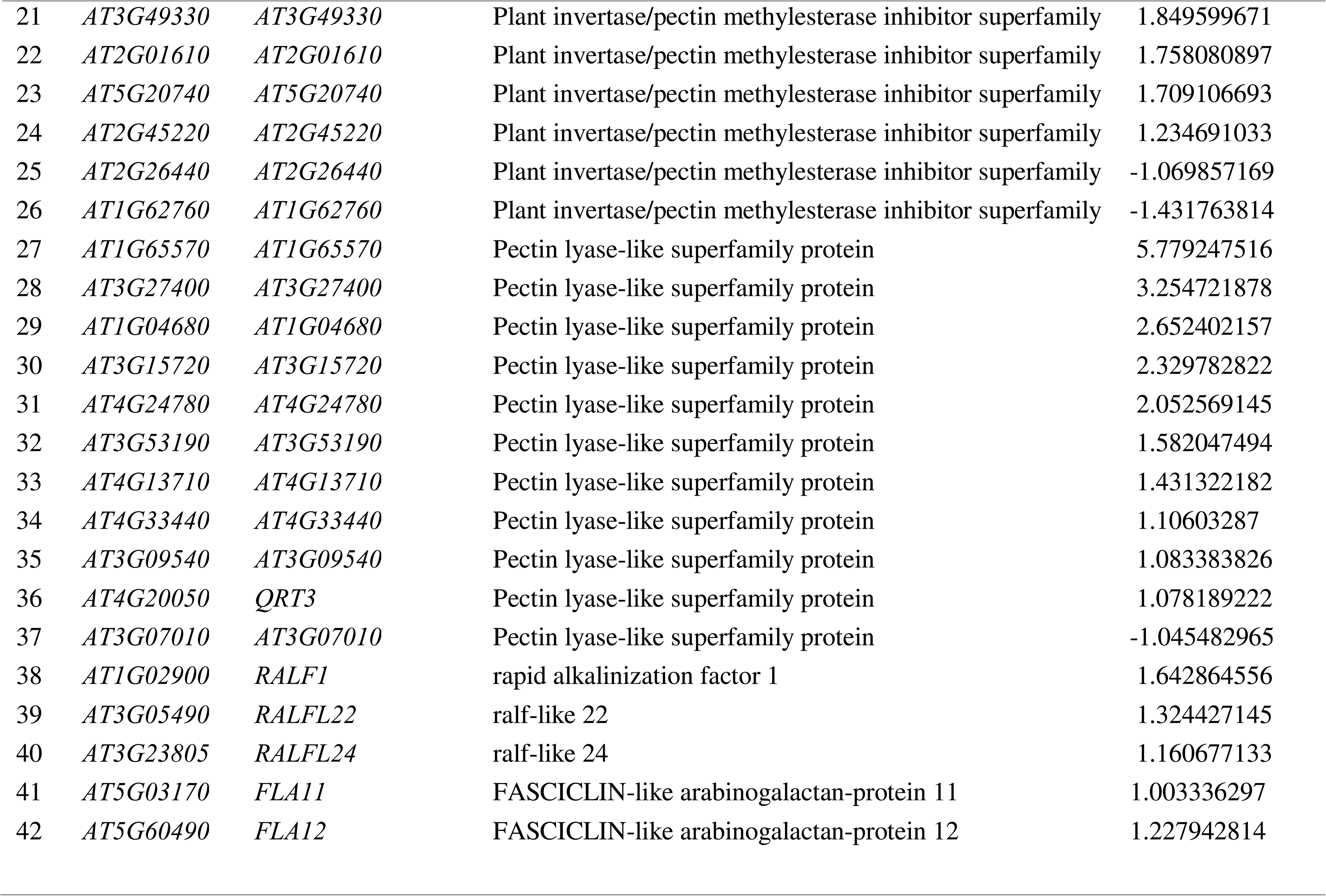
List of differentially expressed genes (DEGs) associated with cell wall acetylation, remodeling and integrity.

The following genes were represented in GO category involved in cell wall remodeling, *XYLOGLUCAN ENDOTRANSGLUCOSYLASE/HYDROLASES* (*XTHs*), *POLYGALACTURONASES* (*PGs*), *POLYGALACTURONASE INHIBITING PROTEINS* (*PGIPs*), *BETA-GALACTOSIDASES* (*BGALs*), *BETA-XYLOSIDASES* (*XYLs*), *GLYCOSYL HYDROLASES* (*GHs), PECTIN METHYL ESTERASES* (*PMEs*), and *PECTATE/PECTIN-LYASES* (*PLs*). XTHs are xyloglucan-specific hydrolases that have endotransglucosylase and/or endohydrolase activity and the identified *XTHs* i.e., *XTH8*, *XTH15*, *XTH30*, and *XTH32* are involved in cell wall extensibility (Rose et al., 2002, Miedes et al., 2011). The pectin-degrading enzyme, *PG2*/*ADPG2* plays a role in floral organ abscission, while *PGIP2* belongs to a class of defense proteins that protect plants by inhibiting fungal polygalacturonases and enhancing pathogen resistance (Ogawa et al., 2009, Ferrari et al., 2003). *BGAL3* and *BGAL7* are identified as β-galactosidases that are involved in pectin remodeling (Moneo-Sanchez et al., 2019, Collins et al., 2019). *XYL4*/*BXL4* is a β-D-xylosidase that specifically acts on xylan oligosaccharides and is also known to modulate systemic immune signalling in Arabidopsis (Minic et al., 2004, Bauer et al., 2022). Several other glycosyl hydrolases, PMEs and PLs are also identified, which might also contribute in remodeling of the cell wall. Some DEGs were found to be associated with cell wall integrity such as *RAPID ALKALINIZATION FACTORS* (*RALFs*) and *FASCICLIN-LIKE ARABINOGALACTAN-PROTEINS* (*FLAs*). The identified *RALFs,* i.e., *RALF1*, *RALF22*, and *RALF24*, are peptide hormones that bind to specific cell wall integrity receptors and modulate stress responses and root growth in Arabidopsis and are involved in plant immunity (Yu et al., 2020, Wang et al., 2020, Liu et al., 2024). The identified *FLAs* i.e., *FLA11* and *FLA12,* are known to be involved in sensing secondary cell wall complexes to finetune the deposition of lignin and cellulose in the cell wall (Ma et al., 2022). All these results suggest the role of MYB46 in different cell wall processes that were not known before.

### ChIP-sequencing identifies MYB46 genomic binding sites and novel cell wall-related targets within the promoter region belonging to cell wall acetylation, remodeling and integrity

To identify genome-wide MYB46 binding sites, we performed chromatin immunoprecipitation followed by sequencing (ChIP-sequencing). For this study, we used 15-day-old seedlings of GFP-tagged stable transgenic lines of *At*MYB46. The samples were prepared using anti-GFP agarose beads in three replicates with a negative control without GFP. The peak annotation files obtained after sequencing were aligned to *Arabidopsis thaliana* reference genome on the Integrative Genomics Viewer (IGV) software (Figure S6). Depending on the number of peaks obtained in ChIP-sequencing, we choose *At*MYB46-GFP-2 (S4_peaks) with the highest number of peaks for further analysis. In this sample, we identified 11806 unique MYB46 binding peaks distributed all over the genome. Of the total, 9% of peaks are present in the promoter region, half of the peaks (50%) are present in the exonic region, 19% in the intronic region, 3% in the intergenic region, and 19% in the transcription termination site (TTS) (Figure 6a). The identified peaks from all the genomic regions primarily correspond to several known MYB46-regulated processes: 55 genes encoding MYB transcription factors, 37 genes involved in lignin biosynthesis pathways, 17 genes associated with cellulose biosynthesis and 9 genes related to xylan biosynthesis. Analysis of this data further revealed additional gene categories under potential direct regulation by MYB46: 57 genes involved in cell wall acetylation, 67 genes associated with cell wall remodeling, and 25 genes related to cell wall integrity maintenance (Figure 6b). This finding might expand our understanding of MYB46’s regulatory network. The regulation of secondary cell wall biosynthesis by MYB46 that activates a suite of genes involved in the biosynthesis of three major cell wall components i.e., cellulose, lignin, and xylan, has been well studied (Kim et al., 2013, Kim et al., 2014). Also, several other MYB transcription factors have been identified as important regulators of secondary cell wall biosynthesis (Ko et al., 2009, Nakano et al., 2015). Therefore, we focused our analysis on gene families involved in polysaccharide acetylation, cell wall remodelling, and integrity whose transcriptional regulation is not known. We specifically analyzed MYB46-associated gene targets within the promoter-TSS (transcription start site) region. We found gene targets belonging to cell wall acetylation such as *AtGELPs* (*AtGELP4, AtGELP12, AtGELP35, AtGELP71, AtGELP77, AtGELP89*) and *TBLs* (*TBL42* and *TBL21*), cell wall remodeling such as *PMEs* (*PME22* and *PME56*), *XTHs* (*XTH13* and *XTH3*), *BGALs* (*BGAL11*), and cell wall integrity such as *RALFs* (*RALF15* and *RALF9*) and *WALL-ASSOCIATED KINASES LIKE* (*WAKL5* and *WAKL18*) (Figure 6c). This further confirmed changes in cell wall composition and polysaccharide acetylation (Figure 2 and Figure 3) could be directly through MYB46 by regulating above mentioned genes.

**Figure 6.**
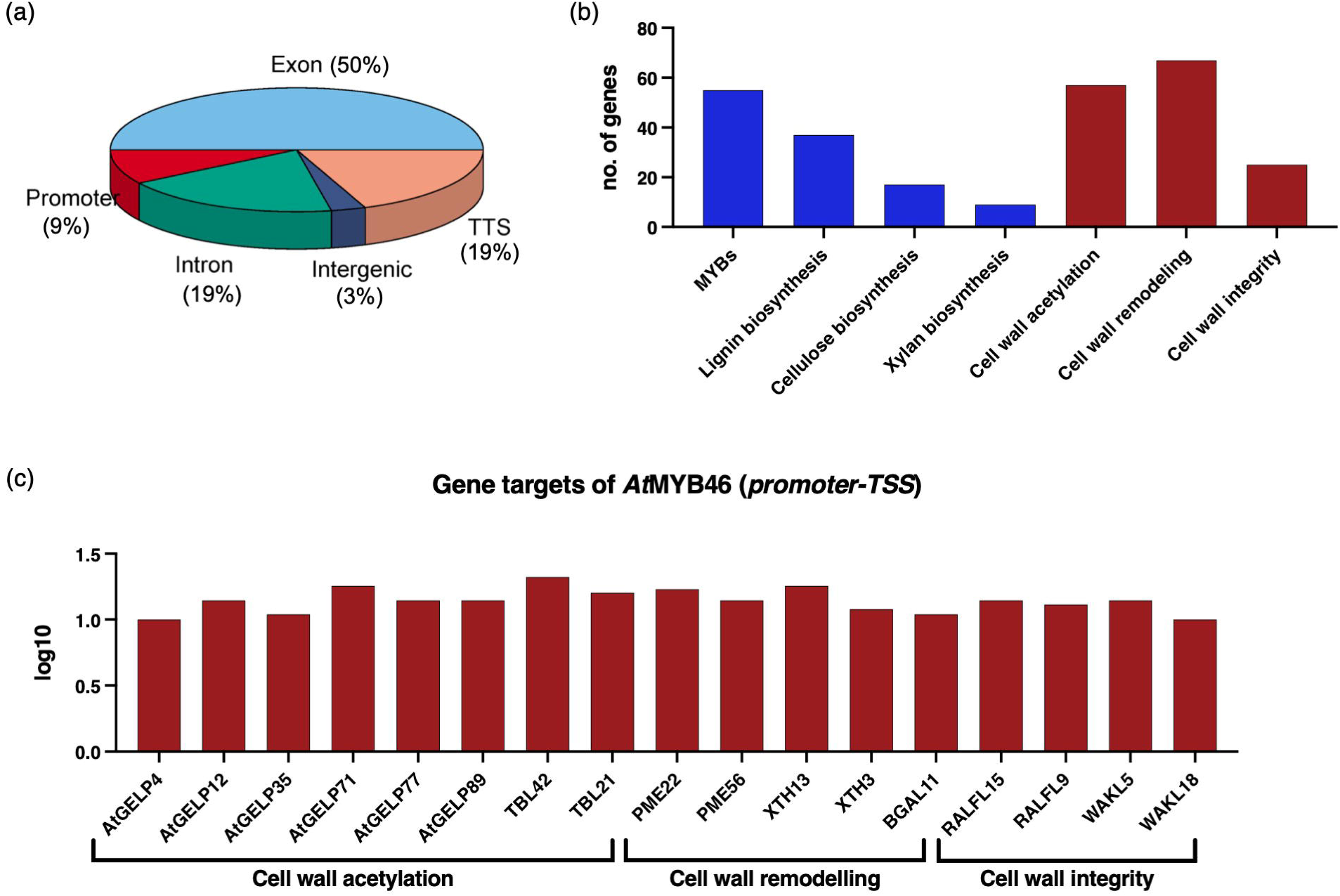
Chromatin immunoprecipitation (ChIP) sequencing of *At*MYB46 reveals novel cell wall-related targets. (a) Pie chart showing the percentage of the *At*MYB46 binding peaks in various genomic regions such as promoter (9%), exon (50%), intron (19%), intergenic (3%), and TTS-transcription termination site (19%) as obtained in ChIP-sequencing data. (b) Graph depicting the number of genes found in ChIP-sequencing data involved in various cell wall-related processes. (c) Representation of some gene targets of *At*MYB46 that are binding specifically in the promoter region and are involved in cell wall acetylation, remodeling and integrity.

### Six hundred differentially expressed genes bound by MYB46 are likely to be novel direct targets of MYB46

To identify more likely direct targets of MYB46, we combined the RNA-sequencing and ChIP-sequencing data sets and found 600 common genes that are differentially expressed upon MYB46 overexpression and also bound by MYB46 (Figure 7a). Pathway analysis of these common 600 genes by GO term represented their involvement in biological processes such as cell wall modification, thickening, organization, and biogenesis (Figure 7b). From this combined data set, we found 19 genes that were associated with cell wall acetylation, remodeling and integrity (Table 2). Six genes involved in cell wall acetylation were identified and these were *AtGELP17*, *TBL33*, *TBL34*, *ACLA-2*, *ACLB-2*, and *REDUCED WALL ACETYLATION* (*RWA3*).

**Figure 7.**
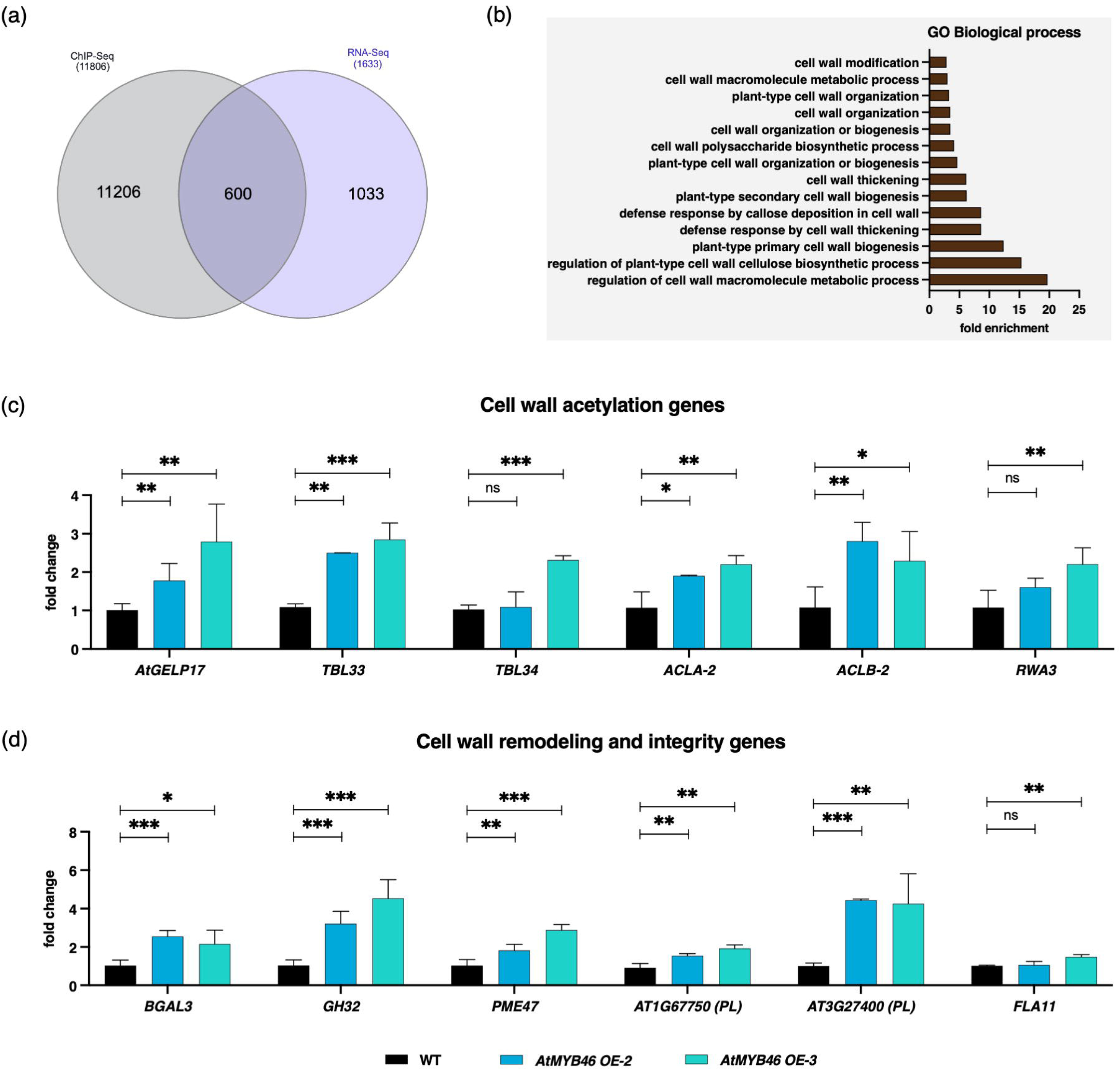
Validation of differentially expressed genes (DEGs) that are bound by MYB46. (a) Venn diagram showing overlapped genes after combining RNA-Seq and ChIP-Seq data. (b) Pathway analysis by Gene Ontology (GO); biological process, of 600 common DEGs that are bound by MYB46 (b). The relative expression of several DEGs also bound by MYB46, involved in cell wall acetylation (c), cell wall remodeling and integrity (d). Data represents mean ± SE, *n* = 3-4 biological replicates, Student’s t-test at \*\*\*\**p* ≤ 0.001, \*\*\**p* ≤ 0.01, \*\**p* ≤ 0.05, * *p* ≤ 0.1.

**Table 2.** List of selected genes found common in RNA-Seq and ChIP-Seq data sets.

| Gene ID | Gene name | Gene description | log2FoldChange | ChIP-Seq |
| --- | --- | --- | --- | --- |
| **Cell wall acetylation** |  |  |  |  |
| *AT1G31550* | *AtGELP17* | GDSL-like Lipase/Acylhydrolase superfamily protein | 1.509190313 | exon |
| *AT2G40320* | *TBL33* | TRICHOME BIREFRINGENCE-LIKE 33 | 1.186715714 | TTS |
| *AT2G38320* | *TBL34* | TRICHOME BIREFRINGENCE-LIKE 34 | 1.902886754 | intron |
| *AT1G60810* | *ACLA-2* | ATP-citrate lyase A-2 | 1.433675387 | intron |
| *AT5G49460* | *ACLB-2* | ATP citrate lyase subunit B 2 | 1.18969785 | intron |
| *AT2G34410* | *RWA3* | O-acetyltransferase family protein | 0.891122667 | exon |
| **Cell wall remodeling and integrity** |  |  |  |  |
| *AT4G36360* | *BGAL3* | beta-galactosidase 3 | 1.616406697 | intron |
| *AT1G45130* | *BGAL5* | beta-galactosidase 5 | 0.983506414 | intron |
| *AT1G68560* | *XYL1* | alpha-xylosidase 1 | 0.964822189 | exon |
| *AT1G64390* | *AtGH9C2* | glycosyl hydrolase 9C2 | 1.612039837 | exon |
| *AT1G62660* | *BFRUCT3* | Glycosyl hydrolases family 32 protein | 2.098290046 | exon |
| *AT2G45220* | *PME17* | pectin methylesterase inhibitor superfamily | 1.234691033 | exon |
| *AT5G04970* | *PME47* | pectin methylesterase inhibitor superfamily | 6.417539094 | exon |
| *AT1G67750* | *AT1G67750* | Pectate lyase family protein | 1.440412285 | promoter |
| *AT1G65570* | *AT1G65570* | Pectin lyase-like superfamily protein | 11.55849503 | exon |
| *AT3G27400* | *AT3G27400* | Pectin lyase-like superfamily protein | 3.254721878 | exon |
| *AT4G33440* | *AT4G33440* | Pectin lyase-like superfamily protein | 1.10603287 | exon |
| *AT3G53190* | *AT3G53190* | Pectin lyase-like superfamily protein | 1.582047494 | intron |
| *AT5G03170* | *FLA11* | FASCICLIN-like arabinogalactan-protein 11 | 1.003336297 | TTS |

Twelve cell wall remodeling genes were identified and these were *BGAL3*, *BGAL5*, *XYL1*, *AtGH9C2*, *BFRUCT3*, *PME17*, *PME47*, and five genes from pectate lyase family and pectin lyase-like superfamily. A single cell wall integrity gene i.e., *FLA11* was identified. Some of these genes were validated by RT-PCR in *At*MYB46 OE-2 and *At*MYB46 OE-3 lines. The expression of cell wall acetylation genes such as *AtGELP17*, *TBL33*, *TBL34*, *ACLA-2*, *ACLB-2*, and *RWA3* were upregulated as found in RNA-Seq data (Figure 7c). *At*GELP17 could be a probable acetyl esterase since it belongs to same clade Id to which *At*GELP7 belongs and might be involved in polysaccharide deacetylation (Rastogi et al., 2022). TBL33 and TBL34 are xylan *O*-acetyltransferases (XOATs) and involved in xylan acetylation (Yuan et al., 2016a, Yuan et al., 2016b). ACLA and ACLB are two subunits of ATP-citrate lyase that generate cytosolic acetyl-CoA pool as a substrate for acetylation pathway (Zhong et al., 2020). RWA3 is specifically expressed in inflorescence stem and regulate xylan acetylation by regulating acetyl donor shuttle between cytosol and Golgi lumen (Lee et al., 2011). All of these genes could be the probable direct targets of MYB46, through which it might be regulating polysaccharide acetylation level in the cell wall. The expression of cell wall remodeling and integrity genes such as *BGAL3*, *BFRUCT3* (*GH32*), *PME47*, two genes i.e., *AT1G67750* from pectate lyase family, *AT3G27400* from pectin lyase-like superfamily and *FLA11* were also upregulated suggesting that these could be the probable MYB46 targets for the regulation of cell wall remodeling and integrity mechanism (Figure 7d). Altogether, this data identified novel probable direct targets of MYB46 through which the transcriptional regulation of polysaccharide acetylation, cell wall remodeling and integrity might be regulated (Figure 8).

**Figure 8.**
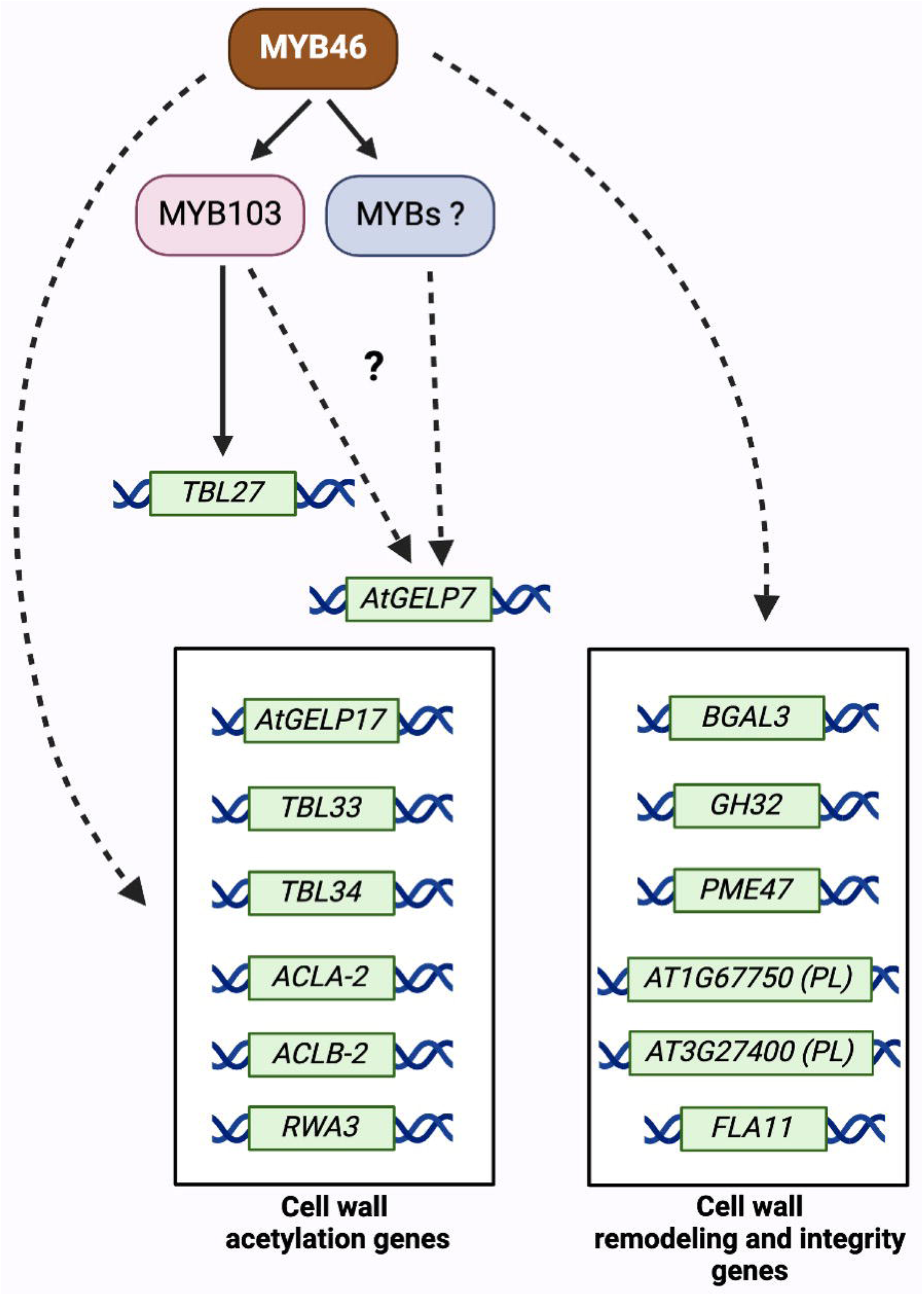
A proposed model showing the transcriptional regulation of cell wall acetylation, remodeling and integrity genes by MYB46.

## Discussion

In Arabidopsis, xylan, xyloglucan, pectin and mannans are *O*-acetylated. The *O*-acetyl substitution patterns, content and specificity to polysaccharides decide its role in plant growth, development and defence. The role of these polysaccharides is mainly explored by characterizing different polysaccharide acetyltransferases and esterases. Although polysaccharide-specific *O*-acetyl transferases from the TBL family are well characterized in Arabidopsis, few polysaccharide-specific acetyl esterases are known. Moreover, a balanced level of *O*-acetyl transferases and esterases might be necessary to balance acetylation both in Golgi and apoplast. But the exact mechanism behind maintaining polysaccharide acetylation is relatively unknown which could be regulated through transcription factors. Therefore, we studied the role of MYB46 in polysaccharide *O*-acetylation using mainly cell wall chemotyping, RNA sequencing and Chip-sequencing analysis.

We became curious to study *At*GELP7 because its expression was very high in MYB46 overexpressing lines (Rastogi et al., 2022) (Figure 4a, b). In fact, out of all the genes, GELP7 was one of the highly expressed ones as compared to cellulose, lignin or xylan biosynthetic genes. We recently identified this *At*GELP7 as acetyl xylan esterase (Rastogi et al., 2022). Therefore, we measured esterase activity in different tissue types of MYB46 overexpression lines and tried to correlate it with increased expression of *At*GELP7. We found a slight increase in the specific esterase activity, particularly in cauline leaf and main stem bottom tissue, which correlated with the higher expression of GELP7 esterase (Figure S3a, b). Moreover, we found a slight alteration in the total cell wall acetyl content upon MYB46 overexpression, but we observed significant changes in the acetylation level on individual polysaccharides such as xylan, xyloglucan and pectin (Figure 3). In the leaf tissue of MYB46 overexpression lines, the xylan acetylation was reduced, and pectin and xyloglucan showed increased acetylation. In stem tissue, particularly dried main stem showed reduced xylan acetylation and increased acetylation on pectin with no change in acetylation level on xyloglucan (Figure 3c, d, e, f). Collectively, these findings indicated that when xylan acetylation decreases, there is a compensatory increase in the acetylation on either pectin or xyloglucan or on both, which helps maintain overall acetylation levels of polysaccharides in the cell wall. Such kind of compensatory effects have been observed in previous studies. Reduced xylan acetylation in poplar seemed to upregulate RWA gene expression as a compensatory mechanism (Ratke et al., 2015, Pawar et al., 2017). Overexpression of *An*AXE1 and *At*AXE1/*At*GELP7 in Arabidopsis led to reduced xylan acetylation, which was compensated by enhanced pectin acetylation (Pawar et al., 2016, Rastogi et al., 2022). Similarly, overexpression of *At*GELP53 led to reduced xyloglucan acetylation, which was counterbalanced by elevated acetylation levels in both pectin and xylan (Rastogi et al., 2024). Despite these observed patterns of compensation, the underlying regulatory mechanisms remain elusive and need further attention.

Transcriptomic studies in MYB46 overexpression plants identified a total of seven differentially expressed *At*GELPs, and six TBLs (Table 1). Among identified *At*GELPs, only *At*GELP7 is characterized as acetyl xylan esterase. Among identified TBLs, most of them were xylan *O*-acetyltransferases. The simultaneous increase in expression of both deacetylating enzymes (xylan-specific acetyl esterase; *At*GELP7) and acetylating enzymes (xylan-specific acetyltransferases; ESK1/TBL29, TBL33, and TBL34) indicates that cell wall polysaccharide acetylation involves sophisticated regulatory mechanisms that maintain precise acetylation homeostasis in the cell wall. However, the observed alterations in polysaccharide acetylation in MYB46 overexpression lines, specifically reduced xylan acetylation, may be primarily attributed to the upregulated expression of *At*GELP7, which deacetylates xylan and thereby reduces xylan acetylation. The increased *At*GELP7 expression observed in MYB46 lines indicates that MYB46 might function as a direct regulator of *At*GELP7. Transactivation assays performed in Nicotiana using the MYB46 transcription factor and *At*GELP7 promoter revealed that MYB46 does not directly bind to or regulate the *At*GELP7 promoter region (Figure S4). Our further studies in GUS stable lines of *At*GELP7 promoter revealed that MYB46 is indirectly regulating the expression of *At*GELP7 gene through downstream regulators (Figure 4d, e). Several MYB transcriptional regulators have been identified as directly regulated by MYB46 (Ko et al., 2009, Nakano et al., 2015). Among them, MYB103 was recently identified to positively regulate cell wall acetylation by directly activating TBL27 under aluminium (Al) stress (Wu et al., 2022). It is possible that MYB46 may be regulating *AtGELP7* gene expression through MYB103 or via other MYB TFs, as shown in our model (Figure 8). This transcriptional regulation by MYB46 via MYB103/MYBs of *AtGELP7* gene needs further investigation. To identify the possible targets that might be more specifically involved in maintaining the acetylation homeostasis, we combined Chip-Seq and RNA-Seq data and identified the following genes belonging to the polysaccharide acetylation pathway: *AtGELP17*, *TBL33*, *TBL34*, *ACLA-2*, *ACLB-2*, and *RWA3*. Among these genes, only *AtGELP17* is uncharacterised. However, its phylogenetic placement in the same clade as *At*GELP7 suggests it may also act as a polysaccharide acetyl esterase. Altogether, these genes could be direct targets of MYB46 through which it might be positively regulating cell wall polysaccharide acetylation. RT-PCR analysis further confirmed that MYB46 positively regulates the expression of these genes (Figure 7c). The direct regulation of these genes by the MYB46 transcription factor needs to be investigated in future through transactivation studies and electrophoretic mobility shift assay (EMSA).

Apart from understanding the transcriptional regulation of polysaccharide acetylation, we were also curious to understand the changes in the cell wall composition upon MYB46 overexpression in different tissue types. The extensive cell wall compositional characterization revealed that lignin content was higher in all different stem tissues, like side stem, main stem bottom, main stem top, and dried main stem and no change in lignin content was observed in leaf tissue such as in rosette leaf, cauline leaf and dried rosette leaf upon MYB46 overexpression, suggesting that MYB46 is positively regulating lignin biosynthesis, particularly in stem tissue (Figure 2a, b). The cellulose content was found higher only in mature leaf and stem tissues, suggesting that MYB46 positively regulates cellulose biosynthesis by accumulating more cellulose in the later stages of plant development (Figure 2c, d). The composition of matrix polysaccharides such as hemicelluloses and pectin were identified by analyzing monosugar composition of cell walls across different tissue types in MYB46 overexpressing lines. Among hemicelluloses, xyloglucan and xylan are the predominant hemicelluloses in the leaf and stem, respectively. Analysis of the leaf tissues showed an increasing trend of hemicellulosic glucose (a part of xyloglucan) from 4% in rosette leaf to 10% in cauline leaf and reaching to 12% in dried rosette leaf. In contrast, the composition of other monosugars like fucose, rhamnose, arabinose, galactose, mannose, xylose, and galacturonic acid was found reduced in MYB46 overexpression plants (Figure 2e, f and S1a). Analysis of stem tissues showed enhanced hemicellulosic xylose content in main stem bottom and dried main stem tissues. In addition, hemicellulosic glucose content was also increased in side stem, main stem bottom and main stem top, whereas other monosugars showed a similar reducing trend in MYB46 overexpressor lines (Figure 2g, h and S1b, c). This data indicates that MYB46 overexpression redirects the metabolic flux towards major cell wall polysaccharides such as cellulose, xyloglucan and xylan while reducing other sugar levels to maintain the metabolic balance.

Interestingly, our transcriptomic data also identified several differentially expressed genes involved in cell wall remodelling and integrity processes. Cell wall remodeling and integrity mechanisms appear to be transcriptionally controlled through changes in the expression of various cell wall-modifying enzymes and proteins identified in Table 1, including XTHs, PGs, PGIPs, BGALs, BXLs, GHs, PMEs, PLs, RALFs and FLAs. Moreover, the ChIP-sequencing of MYB46-GFP seedlings identified 9 promoter-binding gene targets belonging to cell wall remodeling and integrity and these are *PME22*, *PME56*, *XTH13*, *XTH3*, *BGAL11*, *RALF15*, *RALF9*, *WAKL5* and *WAKL18* (Figure 6c). The combined RNA-Seq and ChIP-Seq data further revealed more direct probable targets of MYB46 related to these processes and are listed in Table 2. Some of these genes, such as *BGAL3*, *GH32*, *PME47*, *AT1G67750* (*PL*), *AT3G27400* (*PL*) and *FLA11* were further validated by RT-PCR and found to be upregulated (Figure 7d). These findings suggest that MYB46 may directly regulate the expression of these genes, potentially serving as a positive regulator of cell wall modification and integrity mechanisms in plants.

Overall, this research revealed previously unknown genes likely regulated directly by MYB46 that control cell wall processes like polysaccharide acetylation, cell wall remodeling and integrity as shown in the model (Figure 8). The transcriptional regulation of *At*GELP7 esterase by MYB46 via MYB103/MYBs is also highlighted in the model. These findings provide new avenues for investigating how MYB46 exerts transcriptional control over cell wall-related functions, more specifically *O*-acetylation in different polysaccharides and cell wall types.

## Materials and Methods

### Cloning of *AtMYB46* and *AtGELP7* gene

*AtMYB46* (*AT5G12870*) gene was cloned in pCC0995 (containing constitutive 35S promoter) and pSITE-2CA (containing N-terminal GFP tag) by gateway cloning method as explained in (Rastogi et al., 2022). The different regions *AtGELP7* promoter *containing SMRE sites* (*AT1G28580*) gene was cloned in pKGWFS7 (containing glucuronidase-GUS) and pKGWFL7 (containing luciferase-LUC) by the same approach.

### Generation of stable Arabidopsis transgenic lines

Different expression clones (*35S::AtMYB46*, *35S::GFP-AtMYB46*, *pAtGELP7::GUS*) in Agrobacterium GV3101 strain were cultured and resuspended in a transformation medium consisting of 5% sucrose (GRM3063, HiMedia, India) and 0.05% silwet-77 (PCT1554, HiMedia, India). Further, the floral dip transformation of wild-type *Arabidopsis thaliana* plants was done according to the protocol explained in (Clough and Bent, 1998). The seeds were collected from the dried transformed plants. Successfully transformed plants with *35S::AtMYB46* were selected by spraying glufosinate ammonium (BASTA) (C45520, Sigma-Aldrich, Switzerland), whereas successfully transformed plants with *35S::GFP-AtMYB46* and *pAtGELP7::GUS* were selected on kanamycin antibiotic MS plates.

### RNA isolation and qRT-PCR

The total RNA from different tissues of Arabidopsis plants was isolated by Trizol method (15596018, Invitrogen, Canada) and One ug of RNA was used for cDNA synthesis carried out by iScript cDNA Synthesis kit (1708891, BioRad, USA). *ACTIN2* (*AT3G18780*) and *GAPDH* (*AT1G13440*) were used as reference genes for qRT-PCR. The expression of all the genes was analysed using gene-specific qPCR primers as listed in the Supplementary Table. The relative fold change of all genes was normalised to the reference gene and calculated by the ΔΔ method.

### Subcellular localization in Nicotiana

Agrobacterium carrying *35S::GFP-AtMYB46* was cultured at 28°C for two days. The cells were resuspended in infiltration medium (10 mM MES buffer, 10 mM MgCl_2_, pH - 5.6), OD was set to 0.4, and the suspension was induced with 100 µM acetosyringone (RM9145, HiMedia, India). The induced Agrobacterium suspension was then infiltrated in Nicotiana leaves. For nuclei staining, 4’,6-diamidino-2-phenylindole, dilactate (DAPI) () (D3571, Invitrogen, USA) was used in a concentration of 1 ug/ml and infiltrated just before imaging on 3^rd^ day. Small sections from the infiltrated leaves were visualised under LEICA SP8 confocal microscope with 40X magnification.

### Arabidopsis protoplast isolation

The fully expanded leaves of stable *GFP-AtMYB46* Arabidopsis lines were used for protoplast isolation, which was done according to the Tape-Arabidopsis Sandwich method (Wu et al., 2009). Briefly, the upper epidermal layer of the leaf was affixed with time tape and the lower epidermal layer was peeled off using magic tape. The peeled leaves were transferred to a petri plate containing the enzyme solution consisting of 1% cellulase ‘Onozuka’ R10 (Yakult, Tokyo, Japan), 0.25% macerozyme (PCT1531, HiMedia, India), 0.4M mannitol (GRM024, HiMedia, India), 10mM CaCl_2_, 20mM KCl, 0.1% BSA, 20mM MES (pH 5.7). The protoplasts were released in the solution and were directly observed under a LEICA SP8 confocal microscope in a 40X oil immersion lens.

### Toluidine Blue O staining

Inflorescence stem sections of wild-type and *At*MYB46 OE plants were hand sectioned and stained with 0.02% Toluidine Blue O (T3260, Sigma-Aldrich, USA) solution, and images were captured in a Nikon microscope at 10X and 40X magnification.

### Phloroglucinol-HCl staining

Main stem sections of wild-type and *At*MYB46 OE plants were hand sectioned and stained with 3% phloroglucinol (P3502, Sigma-Aldrich, USA) solution. Preparation of phloroglucinol solution and staining of stem sections was done as explained in (Pradhan Mitra and Loque, 2014). The stained sections were visualised at 10X and 40X magnification in a Nikon microscope.

### Preparation of Alcohol insoluble residue (AIR)

Fresh leaf and stem tissues were crushed in liquid nitrogen to a fine powder and dried plant tissues were grounded by Qiagen TissueLyser II. This powder was incubated with 80% ethanol at 70°C for 30 min and centrifuged at 14000 rpm for 15 min. The pellet was then treated with 70% ethanol, which was followed by treatment with chloroform(1):methanol(1). Lastly, the pellet was washed with acetone and dried in a vacuum desiccator. The dried AIR was further used for cell wall analysis.

### Lignin content

The Acetyl Bromide Soluble Lignin (ABSL) method was followed to quantify lignin in different tissues of the Arabidopsis plant. 1 mg of AIR was weighed and treated with 25% acetyl bromide (135968, Sigma-Aldrich, Mongolia) freshly prepared in acetic acid and incubated at 50°C for 2 h with gentle shaking. 2M NaOH and freshly prepared hydroxylamine hydrochloride (159417, Sigma-Aldrich, India) were used to neutralize the reaction mixture. Finally, the absorbance was taken at 280 nm and the lignin content was quantified and represented as mg per g of AIR (Foster et al., 2010).

### Cellulose content

Updegraff method was used to quantify cellulose in different plant AIR samples (Updegraff, 1969). Briefly, 3 mg AIR was first treated with Updegraff reagent [acetic acid (8): nitric acid (1): water (2)] at 100°C for 30 min to remove hemicellulosic sugars and then the remaining pellet (rich in cellulose) was further hydrolysed with 72% sulfuric acid. Glucose content was measured in the resulting hydrolysate by anthrone assay.

### Monosugar composition by ion chromatography (IC)

2 mg AIR of different leaf and stem tissues were hydrolysed by 1.3M HCl at 100°C for 1 h and subsequently neutralised by 1.3M NaOH. The supernatant was diluted 25 times and transferred to vials. Monosugar composition in these samples was analysed using HPAEC-PAD (High-performance anion-exchange chromatography with pulsed amperometric detection) which was conducted on a Dionex ICS-6000 HPIC system (Thermo Fischer Scientific, USA) equipped with a CarboPac PA1 analytical column (4 X 250 mm) and CarboPac PA1 guard column (4 X 50 mm).

The instrument was operated with eluent A (18 mM NaOH) at a flow rate of 1.1 ml/min and a column temperature of 30°C with increasing amounts of eluent B (200mM NaOH) and eluent C (100 mM NaOH and 150 mM sodium acetate). The gradient profile used for sugar analysis was adopted from (Widmer, 2011). Chromeleon 7.0 software (Thermo Fischer Scientific, USA) was used to analyse chromatograms. Monosugar standards of 10 ppm, 5 ppm and 2.5 ppm concentrations were run as follows-fucose, rhamnose, arabinose, galactose, glucose, mannose, xylose, and galacturonic acid to calculate sugar concentration in samples

### Esterase activity

The total crude proteins were isolated from fresh leaf and stem tissues and quantified by Bradford assay. These protein preparations were incubated with p-nitrophenyl acetate (18432, SRL, India), a universal substrate for esterase activity. The reaction was incubated at 37°C for 2 h, and the absorbance of 4-nitrophenol released was taken at 400 nm. The specific esterase activity was calculated using 4-nitrophenol standard curve and represented in nmol per min per mg of total protein.

### Acetyl content analysis in cell wall

1 mg of AIR sample was incubated with 1M NaOH for de-esterification, subsequently neutralized with 1M HCl, and the acetic acid content was analyzed in supernatant by acetic acid kit (K-ACET, Megazyme, Ireland).

### Sequential extraction of cell wall polysaccharides

For sequential extraction, leaf and stem AIR was first incubated with 50mM ammonium formate (50504, SRL, India) buffer at 37°C for 24 h, and pectin-rich fraction I was collected from the supernatant. The residual pellet was further digested with pectate lyase (E-PCLYAN2, Megazyme, Ireland) at 40°C for 24 h, and pectin-rich fraction II was collected from the supernatant. The remaining xyloglucan and xylan rich pellet was digested with xyloglucanase (E-XEGP, Megazyme, Ireland) at 50°C for 2 days, and xyloglucan-rich fraction was collected from the supernatant. These isolated fractions were lyophilized and redissolved in water for further analysis. The remaining xylan-rich pellet was washed with acetone and dried. Acetic acid content was analyzed in all these fractions by acetic acid kit (K-ACET, Megazyme, Ireland). Galacturonic acid content was analysed in pectin-rich fraction I and II by biphenyl method. Xylose content was analysed by xylose kit (K-XYLOSE, Megazyme, Ireland). These fractions were further hydrolysed with 1.3M HCl and run on Dionex ICS-6000 HPIC system as explained above.

### Transactivation, GUS and luciferase assay

The effector construct (*35S::AtMYB46*) and reporter construct (*pAtGELP7::GUS/LUC*) in Agrobacterium were co-infiltrated in Nicotiana leaves and GUS and luciferase assays were performed after 3^rd^ day post-infiltration. For GUS assay, leaf discs were incubated overnight at 37°C in GUS staining solution prepared by dissolving X-Glucurono sodium salt (MB089, HiMedia, India) in DMSO, 50mM sodium phosphate buffer, 100mM potassium ferricyanide (GRM1034, HiMedia, India), 100mM potassium ferrocyanide trihydrate (GRM1048, HiMedia, India), and 0.1% triton. Ethanol (3):acetic acid (1) solution was used for destaining. For luciferase assay, 2.5mM D-Luciferin (LUCK, Goldbio, USA) in 0.01% tween-20 was infiltrated in the construct-infiltrated leaves and directly observed under Image Quant.

### RNA sequencing

Main stem samples were collected in three biological replicates from wild-type and *At*MYB46 OE plants and total RNA was isolated by Trizol method. These RNA samples were quantified on the Qubit 3.0 Fluorometer (ThermoFisher Scientific) using the QubitTM RNA HS Assay Kit following the manufacturer’s protocol. RNA sequencing was performed using Illumina NovaSeq 6000 by Clevergene Biocorp Private Limited, Bengaluru, India and the obtained transcriptomic data was further analysed. Briefly, the sequence reads were processed to remove adapter sequences and low-quality bases using fastp v 0.20. The trimmed reads were mapped onto indexed *Arabidopsis thaliana* Genome assembly TAIR10.1 (GCF_000001735.4) using STAR aligner. The gene level expression values were obtained as read counts using feature-counts software. For differential expression analysis, the biological replicates were grouped which was carried out using the DESeq2 package after normalizing the data using the relative log expression normalization method. The genes with absolute log2 fold change ≥ 1 and adjusted p-value ≤ 0.05 were considered significant. Principal component analysis (PCA), volcano plot and heat map were generated using the SRplot web tool (Tang et al., 2023). For pathway mapping, ShinyGO (v0.80) tool was used to perform over-representation analysis using the KEGG and Gene ontology: Biological process (GO: BP) database.

### ChIP sequencing

15-day-old *At*MYB46-GFP seedlings were crosslinked by incubating 1.5 g of seedlings in 1% formaldehyde prepared in 1X PBS-phosphate buffer saline (1.3 M NaCl, 30 mM Na_2_HPO_4_, 30 mM NaH_2_PO_4_; pH - 7) for 10 min in a vacuum desiccator and repeated once for 10 min. Formaldehyde crosslinking was stopped by adding 2 M glycine to a final concentration of 0.125 M for 5 min in a vacuum desiccator. The crosslinked seedlings were rinsed twice with 1X PBS, once with water, dried on a paper towel and crushed in liquid nitrogen to make a fine powder. For nuclei isolation, the crushed seedling samples were resuspended in cold nuclei isolation buffer (0.25M sucrose, 15 mM PIPES pH-6.8, 5 mM MgCl_2_, 60 mM KCl, 15 mM NaCl, 1 mM CaCl_2_, 0.9 % triton X-100, and 1 mM PMSF) and kept on ice for 15-30 min for complete homogenisation. This slurry was filtered through four layers of cheesecloth, and the filtrate was centrifuged at 11,000 g for 20 min at 4°C to obtain a white pellet (nuclei). The nuclei pellet was resuspended in cold nuclei lysis buffer (50mM Tris-HCl pH-8, 10mM EDTA, 1% SDS, 1mM PMSF) and sonicated in bioruptor sonication device with the settings 30 sec ON / 30 sec OFF for 90 cycles to shear the chromatin in small fragments (200-800 bp). Sheared chromatin was incubated overnight on a rotating wheel at 4°C with anti-GFP IgG agarose beads (BB-GF002B, BioBharti LifeSciences Pvt.Ltd., India) for immunoprecipitation of protein of interest (*At*MYB46-GFP). Agarose beads (bound with chromatin) were washed once with low salt buffer (150 mM NaCl, 20 mM Tris-HCl pH - 8, 0.2% SDS, 0.5% triton X-100, 2 mM EDTA), followed by high salt buffer (500 mM NaCl, 20 mM Tris-HCl pH-8, 0.2% SDS, 0.5% triton X-100, 2 mM EDTA), LiCl buffer (0.25 M LiCl, 1% sodium deoxycholate, 10mM Tris-HCl pH-8, 1% NP-40, 1 mM EDTA), and twice with TE buffer (1 mM EDTA, 10 mM Tris-HCl pH - 8). Immuno-complexes bound to agarose beads were eluted in elution buffer (0.5% SDS, 0.1M NaHCO_3_) and reverse crosslinked by incubating with 5M NaCl at 65°C for 4 h. After reverse crosslinking, protein was digested by incubating samples with 0.5M EDTA, 1M Tris-HCl pH-6.5, and proteinase K for 1.5 h at 45°C. The remaining DNA was further processed by adding an equal volume of chloroform (24):isoamyl alcohol (1), centrifuged at 13,800 g for 15 min at 4°C, and the supernatant was transferred to fresh tubes. The DNA in the supernatant was precipitated by adding 2.5 volume of 100% EtOH and 1/10 volume of 3M sodium acetate pH-5.2 and incubated overnight at -80°C. The DNA pellet was obtained by centrifugation at 13,800g for 15 min at 4°C. The pellet was washed twice with 70% EtOH, dried at room temperature, and dissolved in TE buffer. These enriched DNA samples were quantified using a Qubit 4.0 fluorometer (Thermofisher #Q33238) using a DNA HS assay kit (Thermofisher #Q32851). The library preparation was carried out using Ultra II DNA lib preparation for Illumina Kit (NEBNext #E7645S/L) and insert size was queried on Tapestation 4150 (Agilent) utilizing high sensitive D1000 screentapes (Agilent # 5067-5582) following manufacturers’ protocol. The sequencing of libraries was performed on the Illumina (Novaseq6000) platform by the Neuberg Centre for Genomic Medicine. The peak annotation files obtained after sequencing were aligned to the *Arabidopsis thaliana* reference genome on the Integrative Genomics Viewer (IGV) software.

## Data Statement

The Raw data is submitted to Indian Biological Data Centre (IBDC-INDA accession: INRP000257) and the same is available on INSDC (NCBI/ENA/DDBJ) with BioProject ID PRJEB83879.

## Author Contribution

LR designed and performed all of the experiments. SD helped in analysing ChIP-sequencing data. PP conceptualized, designed, and secured funding for the project. LR and PP wrote the manuscript. All authors read and agree to publish the manuscript

## Acknowledgements

This work was supported by RCB core, SERB-SRG Grant to PP. We would like to thank RCB-Advanced Technology Platform Centre and RCB-Central Instrumentation Facility for access to instruments.

## Conflict of interest

Authors declare no conflict of interest

## Supplementary material

**Supplementary Figure S1.** Monosugar composition analysis (mol%) in cauline leaf, side stem, and main stem (top), analyzed by ion chromatography.

**Supplementary Figure S2.** Total cell wall acetyl content in different leaf and stem tissues in wild-type Arabidopsis plant.

**Supplementary Figure S3.** Esterase activity in different leaf and stem tissues of *At*MYB46 overexpressing lines.

**Supplementary Figure S4.** Transactivation studies in Nicotiana leaves.

**Supplementary Figure S5.** Heat map showing top 30 differentially expressed genes (DEGs) in wild-type and *At*MYB46 overexpression lines found in RNA-sequencing.

**Supplementary Figure S6.** Alignment of peak annotation files obtained after ChIP-sequencing with *Arabidopsis thaliana* reference genome on the Integrative Genomics Viewer (IGV) software.

**Supplementary Table S1.** Monosaccharide composition in sequentially extracted polysaccharide fractions from rosette leaf (a), main stem (bottom) (b), and main stem (dried) of wild-type and *At*MYB46 overexpression (OE) lines in ug/ml, analyzed by ion-chromatography (IC).

**Supplementary Table S2.** Secondary wall MYB-responsive elements (SMRE) sites in promoter sequences of all the gene members of clade Id of Arabidopsis GDSL esterase/lipase protein (GELP) family.

**Supplementary Table S3.** Primer list.

## Notes

### Competing Interest Statement

The authors have declared no competing interest.

