## Supplementary figures and images for "The role of MYB46 in modulating polysaccharide acetylation by mediating the transcriptional regulation of cell wall-related genes in Arabidopsis"

### Supplementary Figure 1

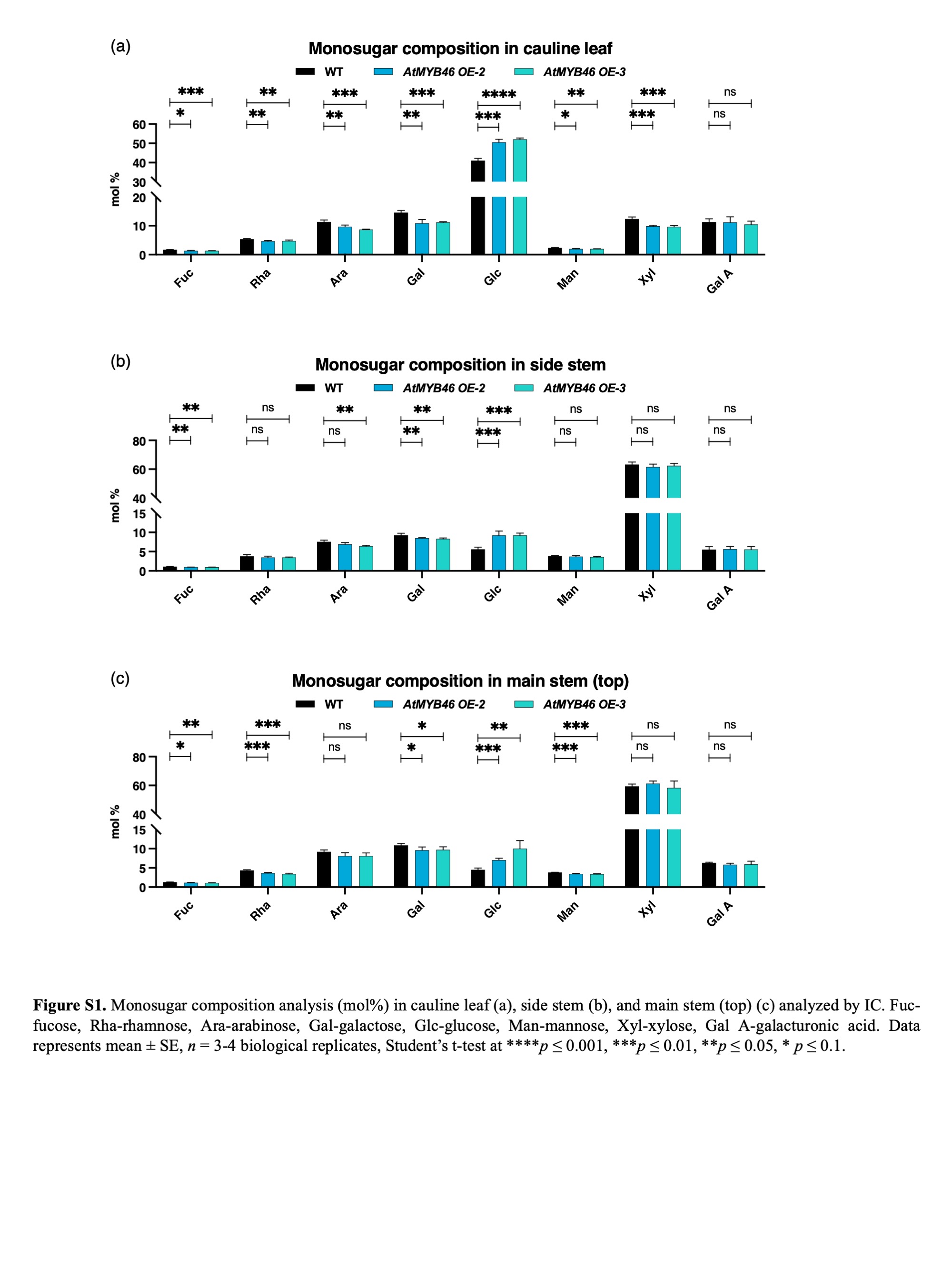

### Supplementary Figure 2

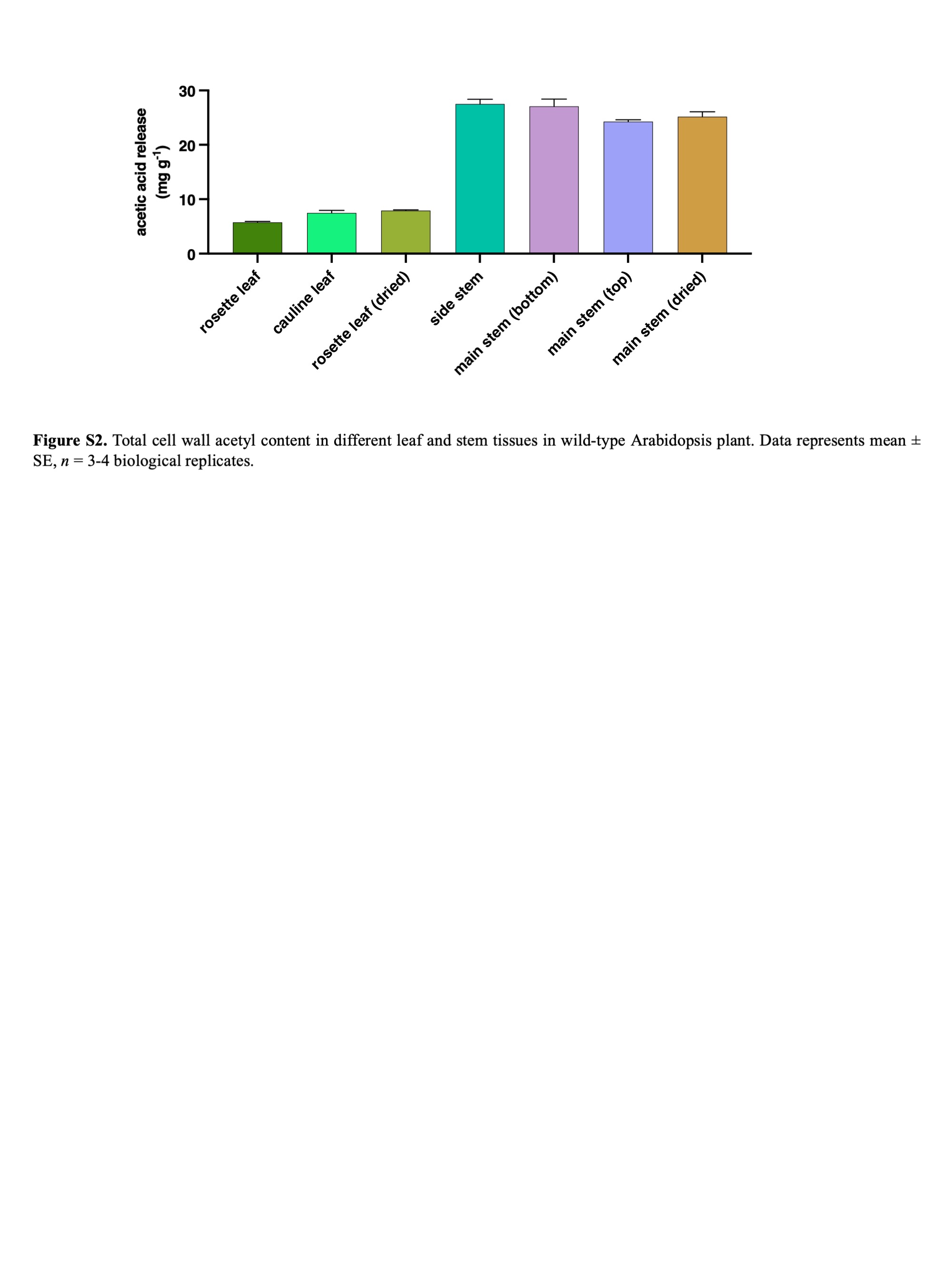

### Supplementary Figure 3

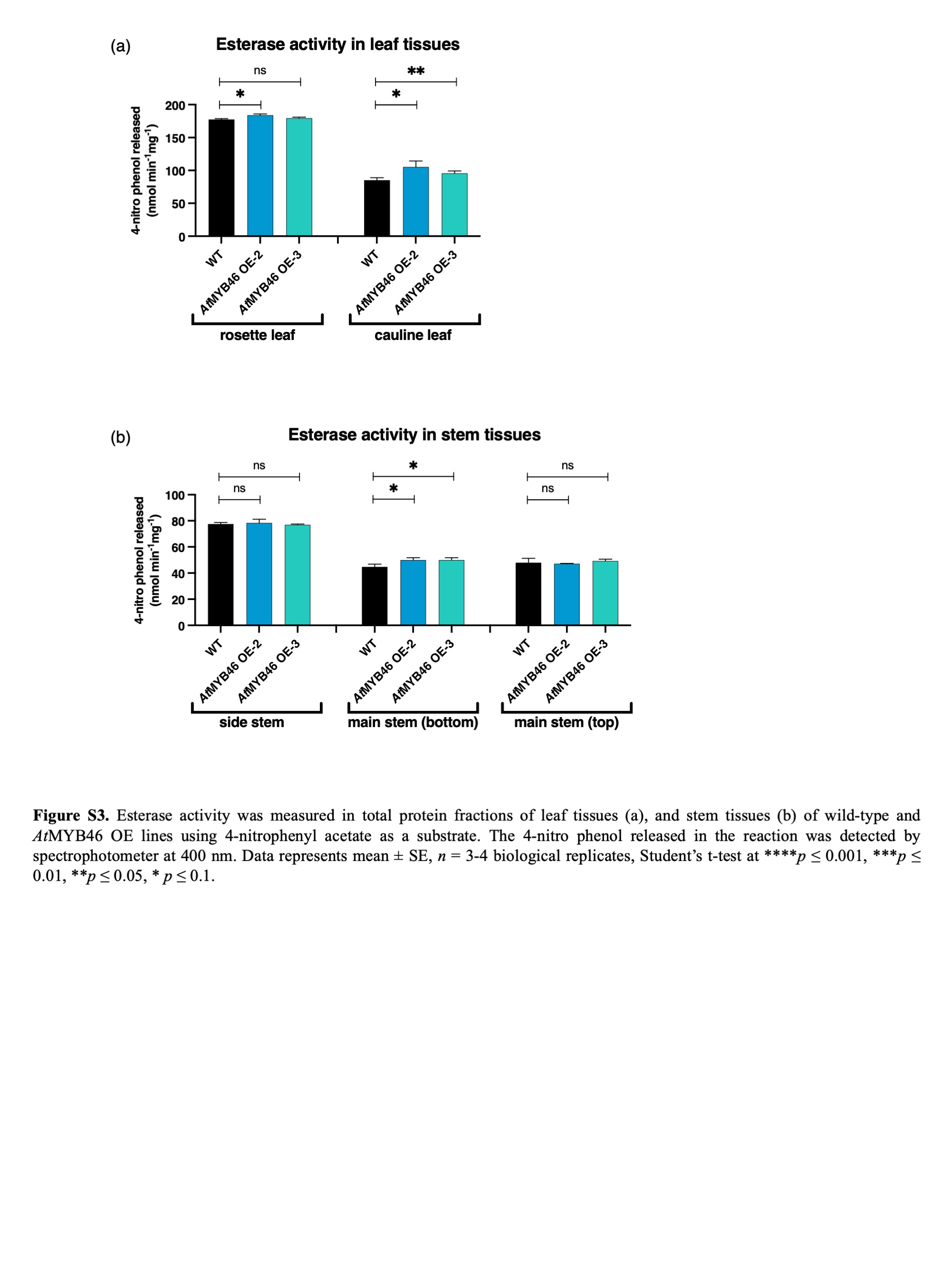

### Supplementary Figure 4

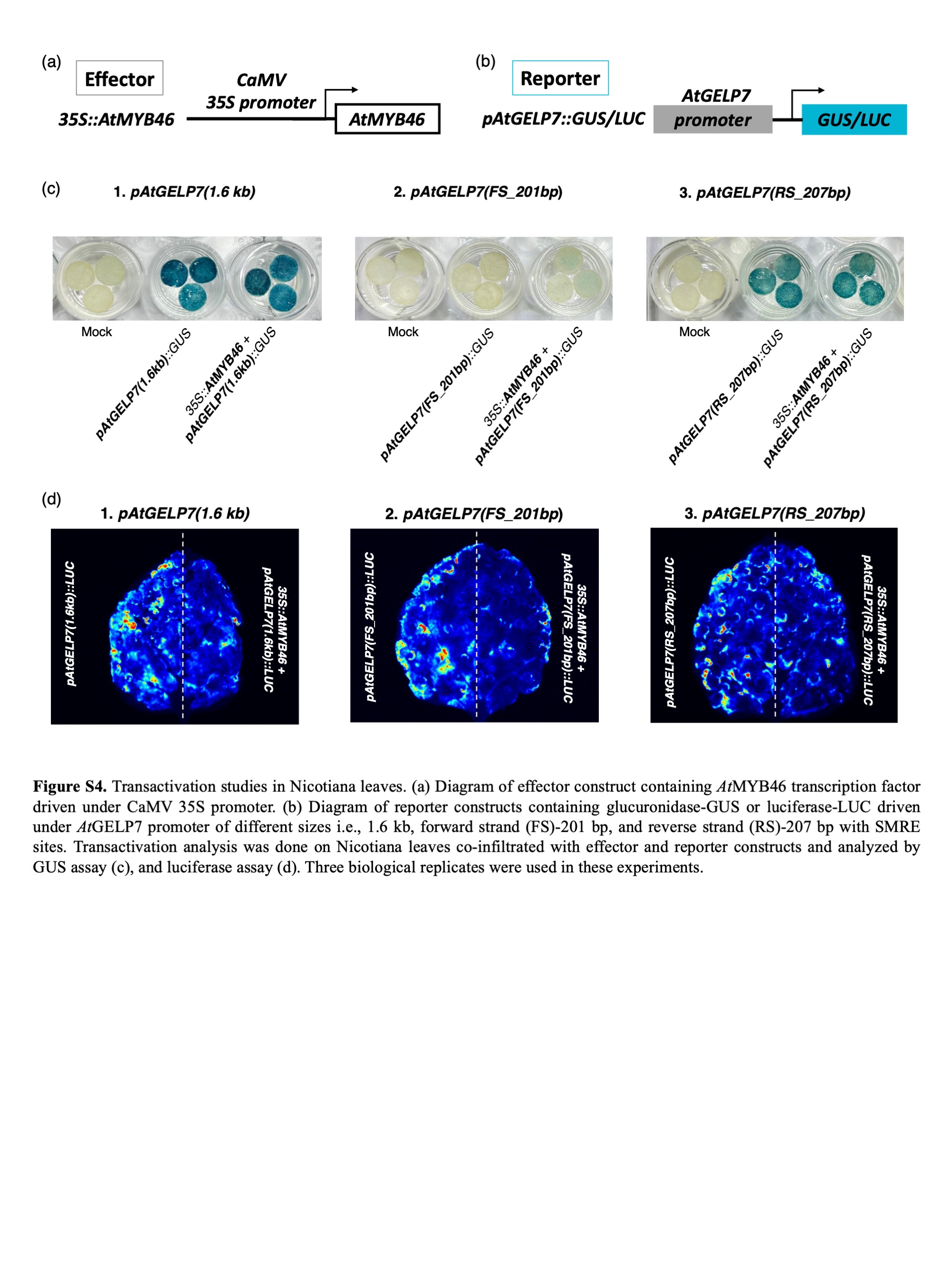

### Supplementary Figure 5

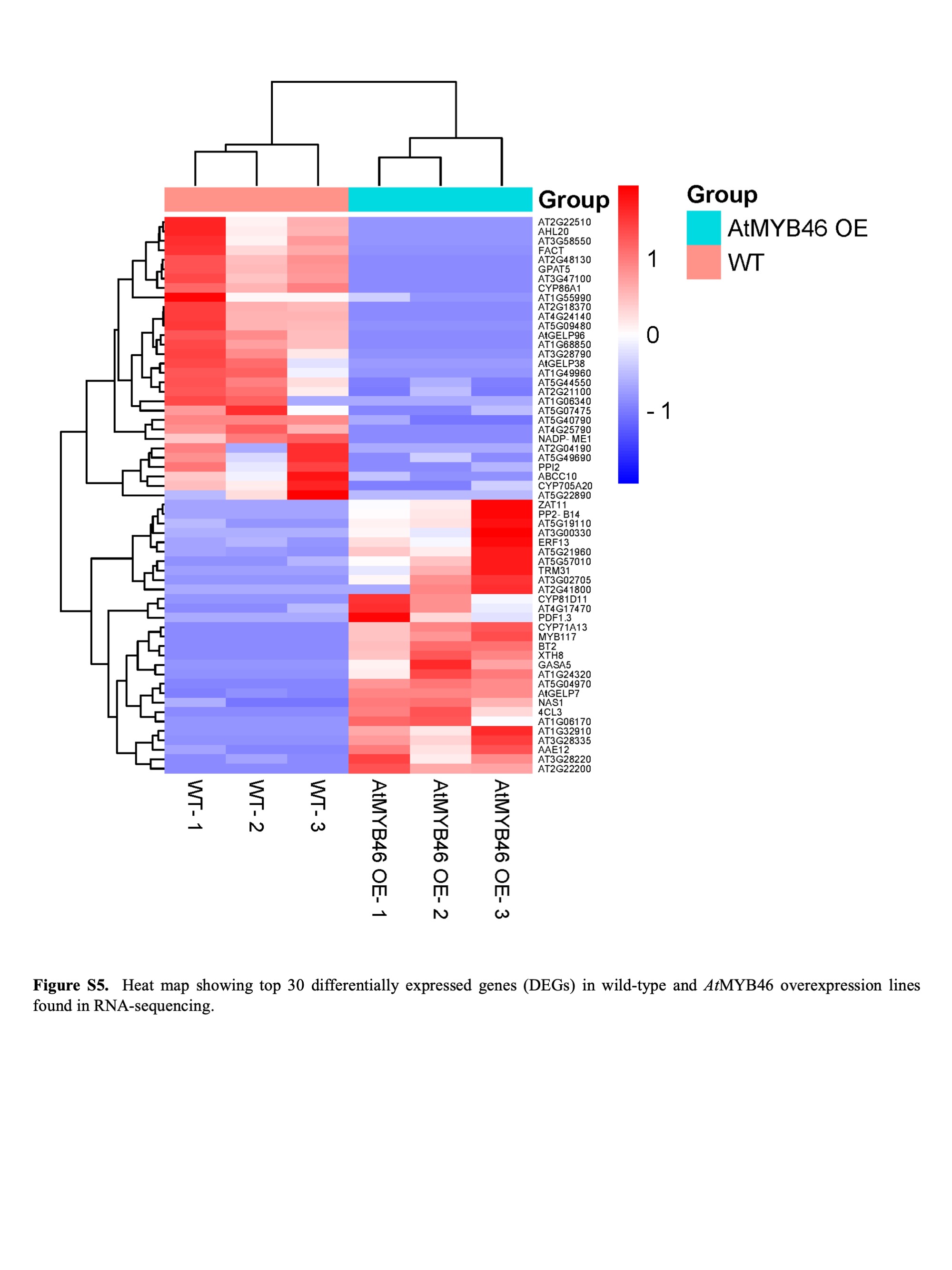

### Supplementary Figure 6

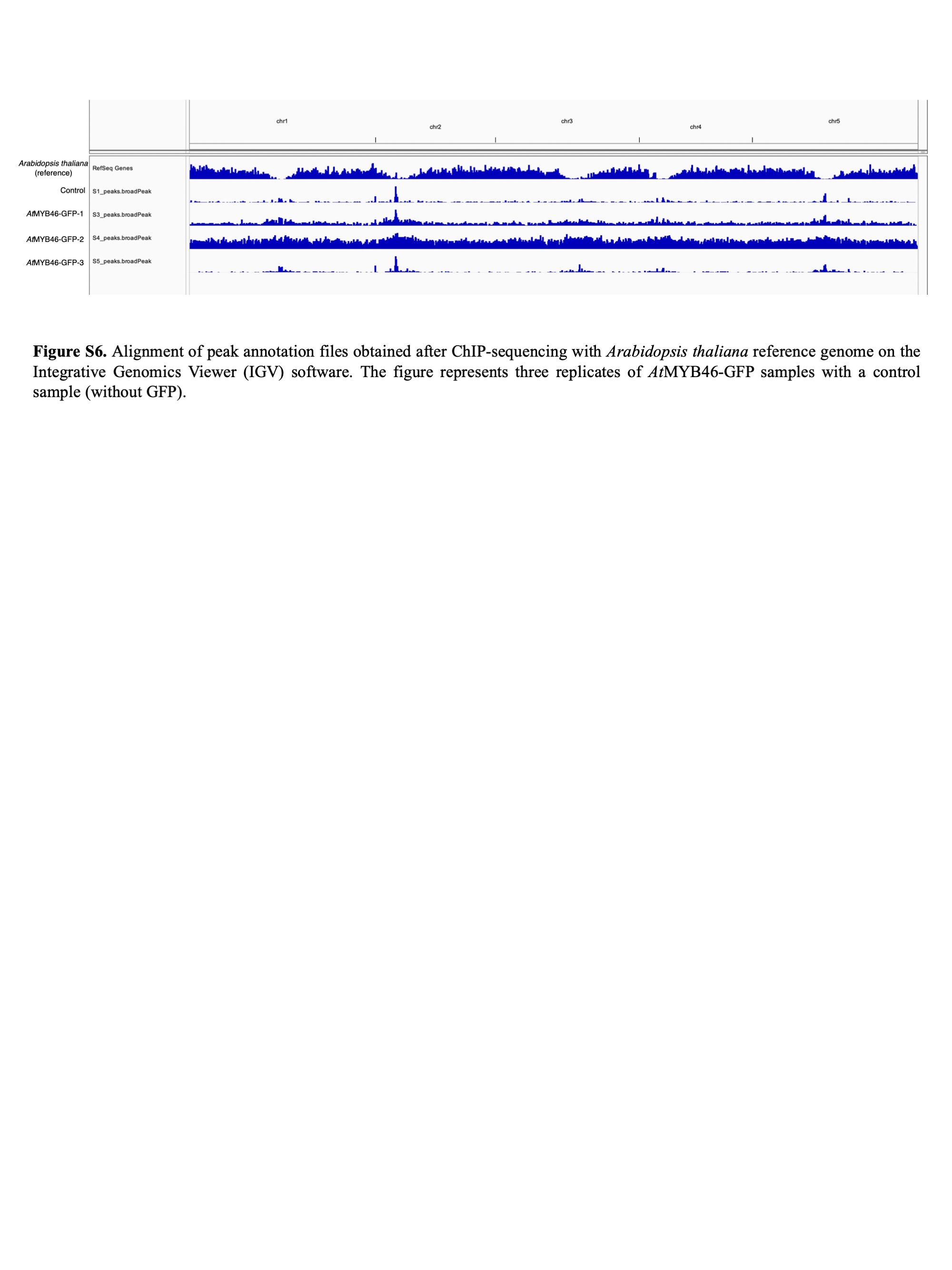
