## Supplementary Tables for "The role of MYB46 in modulating polysaccharide acetylation by mediating the transcriptional regulation of cell wall-related genes in Arabidopsis"

**Table S1.** Monosaccharide composition in sequentially extracted polysaccharide fractions from rosette leaf (a), main stem (bottom) (b), and main stem (dried) of wild-type and *At*MYB46 overexpression (OE) lines in ug/ml, analyzed by ion-chromatography (IC).

(a)

| Fractions | rosette leaf | Fuc | Rha | Ara | Gal | Glc | Man | Xyl | Gal A |
| --- | --- | --- | --- | --- | --- | --- | --- | --- | --- |
| Pectin-rich fraction-I | WT | 10.716 | 11.384 | 62.318 | 108.259 | 8.211 | nd | 10.797 | nd |
| Pectin-rich fraction-I | *At*MYB46 OE-2 | 12.092 | 22.469 | 68.738 | 103.969 | 6.693 | nd | 12.633 | 20.733 |
| Pectin-rich fraction-I | *At*MYB46 OE-3 | 23.646 | 30.899 | 89.838 | 148.372 | 42.015 | nd | 22.325 | 10.723 |
| Pectin-rich fraction-II | WT | 101.984 | 405.101 | 218.386 | 305.526 | 424.234 | 119.548 | 146.898 | 221.987 |
| Pectin-rich fraction-II | *At*MYB46 OE-2 | 86.922 | 389.576 | 198.861 | 291.792 | 222.841 | 104.679 | 163.765 | 212.018 |
| Pectin-rich fraction-II | *At*MYB46 OE-3 | 26.513 | 390.442 | 134.628 | 158.345 | 124.356 | 79.711 | 87.575 | 486.115 |
| Xyloglucan-rich fraction | WT | 25.074 | 76.399 | 63.027 | 74.685 | 163.180 | nd | 38.230 | nd |
| Xyloglucan-rich fraction | *At*MYB46 OE-2 | 49.506 | 62.214 | 66.219 | 82.298 | 144.069 | nd | 69.365 | nd |
| Xyloglucan-rich fraction | *At*MYB46 OE-3 | 64.936 | 60.037 | 60.223 | 86.545 | 180.040 | nd | 126.385 | nd |
| Xylan-rich fraction | WT | 69.284 | 59.614 | 64.485 | 87.121 | 426.079 | 110.791 | 173.261 | nd |
| Xylan-rich fraction | *At*MYB46 OE-2 | 70.081 | 50.219 | 56.373 | 71.639 | 488.120 | 107.187 | 172.850 | nd |
| Xylan-rich fraction | *At*MYB46 OE-3 | 66.183 | 58.452 | 64.743 | 87.364 | 422.025 | 105.855 | 163.353 | nd |

(b)

| Fractions | main stem (bottom) | Fuc | Rha | Ara | Gal | Glc | Man | Xyl | Gal A |
| --- | --- | --- | --- | --- | --- | --- | --- | --- | --- |
| Pectin-rich fraction-I | WT | 22.875 | 62.255 | 170.458 | 281.641 | 127.737 | 120.968 | 96.797 | 158.080 |
| Pectin-rich fraction-I | *At*MYB46 OE-2 | 19.244 | 59.705 | 145.964 | 269.889 | 193.866 | 102.982 | 91.573 | nd |
| Pectin-rich fraction-I | *At*MYB46 OE-3 | 25.791 | 60.043 | 155.601 | 278.556 | 170.488 | 101.528 | 95.306 | 116.134 |
| Pectin-rich fraction-II | WT | 76.891 | 301.830 | 413.311 | 326.911 | 293.502 | 177.697 | 51.763 | 407.237 |
| Pectin-rich fraction-II | *At*MYB46 OE-2 | 307.813 | 598.683 | 963.390 | 657.723 | 615.531 | 396.579 | 370.100 | 444.487 |
| Pectin-rich fraction-II | *At*MYB46 OE-3 | 162.519 | 315.534 | 474.051 | 340.343 | 344.572 | 210.072 | 194.770 | 171.687 |
| Xyloglucan-rich fraction | WT | 69.849 | 36.927 | 80.555 | 84.683 | 96.941 | nd | 73.447 | nd |
| Xyloglucan-rich fraction | *At*MYB46 OE-2 | 44.352 | 31.058 | 69.999 | 67.454 | 99.172 | nd | 55.705 | nd |
| Xyloglucan-rich fraction | *At*MYB46 OE-3 | 45.644 | 29.175 | 61.323 | 64.010 | 86.150 | nd | 55.840 | nd |
| Xylan-rich fraction | WT | 39.013 | 73.605 | 59.372 | 106.041 | 89.521 | nd | 625.923 | nd |
| Xylan-rich fraction | *At*MYB46 OE-2 | 28.516 | 65.426 | 43.638 | 84.812 | 110.519 | nd | 561.214 | nd |
| Xylan-rich fraction | *At*MYB46 OE-3 | 28.066 | 70.784 | 41.907 | 87.188 | 96.619 | nd | 651.561 | nd |

(c)

| Fractions | main stem (dried) | Fuc | Rha | Ara | Gal | Glc | Man | Xyl | Gal A |
| --- | --- | --- | --- | --- | --- | --- | --- | --- | --- |
| Pectin-rich fraction-I | WT | 105.584 | 134.366 | 255.414 | 327.338 | 149.268 | 187.841 | 153.333 | 283.027 |
| Pectin-rich fraction-I | *At*MYB46 OE-2 | 72.227 | 96.011 | 181.658 | 227.925 | 106.024 | 132.754 | 62.485 | 175.116 |
| Pectin-rich fraction-I | *At*MYB46 OE-3 | 96.705 | 163.340 | 232.133 | 301.968 | 168.150 | 191.778 | 119.613 | 303.932 |
| Pectin-rich fraction-II | WT | 119.009 | 298.229 | 357.293 | 329.232 | 236.158 | 237.765 | 84.286 | 567.486 |
| Pectin-rich fraction-II | *At*MYB46 OE-2 | 77.765 | 234.749 | 332.516 | 286.214 | 225.317 | 184.763 | 67.841 | 589.726 |
| Pectin-rich fraction-II | *At*MYB46 OE-3 | 107.554 | 294.022 | 312.220 | 292.322 | 275.600 | 216.460 | 98.294 | 595.004 |
| Xyloglucan-rich fraction | WT | 86.573 | 43.210 | 63.734 | 83.219 | 145.190 | 53.670 | 81.214 | nd |
| Xyloglucan-rich fraction | *At*MYB46 OE-2 | 75.934 | 33.746 | 63.541 | 80.016 | 148.171 | 48.366 | 79.060 | nd |
| Xyloglucan-rich fraction | *At*MYB46 OE-3 | 79.500 | 48.867 | 55.849 | 78.402 | 207.559 | 59.462 | 89.032 | nd |
| Xylan-rich fraction | WT | 50.741 | 76.267 | 59.678 | 97.994 | 171.677 | 130.406 | 523.308 | nd |
| Xylan-rich fraction | *At*MYB46 OE-2 | 48.071 | 67.970 | 61.008 | 101.035 | 147.164 | 118.849 | 512.440 | nd |
| Xylan-rich fraction | *At*MYB46 OE-3 | 53.548 | 82.818 | 59.672 | 107.286 | 207.666 | 151.485 | 622.606 | nd |

**Table S2.** Secondary wall MYB-responsive elements (SMRE) sites in promoter sequences of all the gene members of clade Id of Arabidopsis GDSL esterase/lipase protein (GELP) family. *At*GELP7 is highlighted in blue color.

|  | Clade Id | GELP Ids | Base pairs from promoter | SMRE CONSENSUS  ACC(A/T)A(A/C)(T/C) |
| --- | --- | --- | --- | --- |
| 1. | *AT1G28570* | *AtGELP6* | -1070  -566 | ACCAACT (+) (SMRE2)  ACCAAAC (+) (SMRE3) |
| 2. | *AT1G28580* | *AtGELP7* | -1541  -903 | ACCAACC (-) (SMRE4)  ACCTAAT (+) (SMRE5) |
| 3. | *AT1G28590* | *AtGELP8* | -1141  -41 | ACCAAAT (-) (SMRE1)  ACCAAAT (+) (SMRE1) |
| 4. | *AT1G28600* | *AtGELP9* | -1054  -379  -189  -108 | ACCAAAC (-) (SMRE3)  ACCAAAC (-) (SMRE3)  ACCAAAT (+) (SMRE1)  ACCAACT (+) (SMRE2) |
| 5. | *AT1G31550* | *AtGELP17* | -967  -786 | ACCTAAT (-) (SMRE5)  ACCTAAT (-) (SMRE5) |
| 6. | *AT1G28610* | *AtGELP10* | -1295  -1272  -623 | ACCTACC (-) (SMRE8)  ACCAAAT (-) (SMRE1)  ACCAACC (+) (SMRE4) |
| 7. | *AT2G27360* | *AtGELP53* | -1515  -723 | ACCAAAT (+) (SMRE1)  ACCTAAT (-) (SMRE5) |
| 8. | *AT1G28640* | *AtGELP11* | -188 | ACCTAAC (-) (SMRE7) |
| 9. | *AT1G28660* | *AtGELP13* |  | 0 |
| 10. | *AT1G28670* | *AtGELP14* | -1450  -829  -299 | ACCAAAT (-) (SMRE1)  ACCAACC (-) (SMRE4)  ACCTAAC (+) (SMRE7) |
| 11. | *AT1G28650* | *AtGELP12* | -113  -5 | ACCAAAC (-) (SMRE3)  ACCAAAC (-) (SMRE3) |
| 12. | *AT5G45910* | *AtGELP101* | -247 | ACCAAAT (+) (SMRE1) |
| 13. | *AT3G48460* | *AtGELP72* | -1159 | ACCTACT (+) (SMRE6) |

**Table S3. Primer list.** F; Forward primer, R; Reverse primer

| Oligo name | Target gene ID | Name of gene | Sequence (5´-3´) |
| --- | --- | --- | --- |
| PCWL-395 F | *AT3G53260.1* | *PAL2* | TGTCCAACGGTGAGACTGAG |
| PCWL-396 R | *AT3G53260.1* | *PAL2* | CAAGCTCTTCCCTCACGAAC |
| PCWL-383 F | *AT4G36220.1* | *F5H* | ATGATGGGGATGTTGTCGAT |
| PCWL-384 R | *AT4G36220.1* | *F5H* | ACTCCGTTAAGGCCCACTCT |
| PCWL-776 F | *AT2G30490.1* | *C4H* | ACCGAGCCTGATCTTCACAA |
| PCWL-777 R | *AT2G30490.1* | *C4H* | GGTTGTTTGCTAGCCACCAA |
| PCWL-406 F | *AT1G10670* | *ACLA1* | AAAAAGCAGAGCCCTTGTCA |
| PCWL-407 R | *AT1G10670* | *ACLA1* | GGCTCGCATTTTAGCAAGTC |
| PCWL-408 F | *AT1G60810* | *ACLA2* | GTAGCTGGAGGAGGTGCAAG |
| PCWL-409 R | *AT1G60810* | *ACLA2* | TGAAGGTAGCAGCAACATCG |
| PCWL-410 F | *AT1G09430* | *ACLA3* | GATGACACTGCTGCCTTCAA |
| PCWL-411 R | *AT1G09430* | *ACLA3* | AGCTACCATCGTCCAAATGC |
| PCWL-500 F | *AT1G31550.2* | *AtGELP17* | CTCAGCAACTGCATTGGAAA |
| PCWL-501 R | *AT1G31550.2* | *AtGELP17* | AAGCTCTTTGACCTCGTCCA |
| PCWL-1118 F | *AT2G40320.1* | *TBL33* | GCATCTTTGGACAGCTCGAG |
| PCWL-1119 R | *AT2G40320.1* | *TBL33* | CCTCATACAGTGGTCGGGAG |
| PCWL-1120 F | *AT2G38320.1* | *TBL34* | CCTCAAAGCCACGGTGTTAC |
| PCWL-1121 R | *AT2G38320.1* | *TBL34* | AACCTGGGAAGATCACACGT |
| PCWL-1122 F | *AT5G49460.1* | *ACLB-2* | TGCTATTGGTGGAGACGTGT |
| PCWL-1123 R | *AT5G49460.1* | *ACLB-2* | CAAGTTCCACTGACCCAAGC |
| PCWL-149 F | *AT2G34410* | *RWA3* | TCAGCCATGACATCCTTGAA |
| PCWL-150 R | *AT2G34410* | *RWA3* | CAGACGTAGGCAGCAATGAA |
| PCWL-1124 F | *AT4G36360.1* | *BGAL3* | GCATTGCATGGGTTGAGTCA |
| PCWL-1125 R | *AT4G36360.1* | *BGAL3* | CCTTCGGGCGCATCAAAATA |
| PCWL-1128 F | *AT1G62660.1* | *GH32* | GGGTGACATTGTTTGGGGTC |
| PCWL-1129 R | *AT1G62660.1* | *GH32* | ACCGGTGTAGAGCATGACAA |
| PCWL-1132 F | *AT5G04970.1* | *PME47* | GCCGCACCTAATCACACATT |
| PCWL-1133 R | *AT5G04970.1* | *PME47* | GGAATGTCACATCAACCGCA |
| PCWL-1130 F | *AT1G67750.1* | *Pectate lyase (PL) family* | AGGAATCTAGGCGTGCTCTC |
| PCWL-1131 R | *AT1G67750.1* | *Pectate lyase (PL) family* | GGGTTCCGGGTTTAGGGTTA |
| PCWL-1136 F | *AT3G27400.1* | *Pectate lyase (PL) family* | TGCTCCTAACACCCGCTTTA |
| PCWL-1137 R | *AT3G27400.1* | *Pectate lyase (PL) family* | CAACTCAAAGTGCCTGCTGT |
| PCWL-1138 F | *AT5G03170.1* | *FLA11* | CACCGTGGCTAACTCTGTCT |
| PCWL-1139 R | *AT5G03170.1* | *FLA11* | ATCCCAAACCCGAATCCAGT |
| PCWL-498 F | *AT1G28580* | *AtGELP7 promoter (1.6kb)* | AAAAAGCAGGCTCAAACGCTTTGTTGCGCCTTTT |
| PCWL-499 R | *AT1G28580* | *AtGELP7 promoter*  *(1.6kb)* | AGAAAGCTGGGTAGGAGCTTCATCAAGATAGGC |
| PCWL-739 F | *AT1G28580* | *AtGELP7 promoter (201bp) Forward strand* | CACCGTTGATGTACACATTCATCA |
| PCWL-740 R | *AT1G28580* | *AtGELP7 promoter (201bp) Forward strand* | CTAAGTCTAAAGAAATAGTT |
| PCWL-741 F | *AT1G28580* | *AtGELP7 promoter (207bp) Reverse strand* | CACCGTGTGTGTCAAACCATACTG |
| PCWL-497 R | *AT1G28580* | *AtGELP7 promoter (207bp) Reverse strand* | GGAGCTTCATCAAGATAGGC |
